# Citizen science and detailed searches reveal widespread occurrence of hybrid-like Syrian x Great-spotted woodpeckers

**DOI:** 10.64898/2025.12.22.695873

**Authors:** Antonii Bakai, Tomasz Figarski, Emilia Grzędzicka, Bartłomiej Kusal, Gerard Gorman, Martin Černý, Alena Fišerová, Petr Veselý, Piotr Zielinski, Jerzy Smykla, Zuzanna Pochłopień, Peter Lešo, Krzysztof Czajowski, Maciej Aleksandrowicz, Marcin Borowik, Stanisław Broński, Tomasz Chodkiewicz, Aleksandra Janiszewska, Joanna Kawka, Piotr Kłonowski, Mikołaj Krzyżanowski, Sławomir Kuczmarski, Władysław Lasoń, Kateřina Loudová, Marcin Matysek, Szymon Mazgaj, Iryna Miedviedieva, Beata Murzyn, Mateusz Niedziółka, Peter Puchala, András Schmidt, Tadeusz Sobus, Karolina Tchoń, Ottó Veszelinov, Kristýna Wehrichová, Vadym Zhulenko, Tomasz Ziółkowski, Antoni Życki, Łukasz Kajtoch

## Abstract

The Syrian Woodpecker (hereafter SW) expansion into Europe since the end of the 19th century has enabled its hybridisation with the Great spotted Woodpecker (hereafter GW). This phenomenon, however, has generally been considered sporadic. Here, we examined the occurrence of mixed pairs and hybrids based on our own field observations and citizen science data. Interspecific pairs and individuals of SW and GW were found to be generally scarce but occurred in all parts of the area investigated, not only at the edges of the SW’s range. The proportion of hybrid occurrence in woodpecker populations was found to be similar across the whole SW’s range in Europe and in both urban and rural populations. However, on a local level, hybrids seemed to be more abundant in certain regions, for example, E Austria, Moravia, SE Poland, W Russia, S Ukraine, Bulgaria, E Greece and W Turkey. Generally, females were 1.4 times more frequent among hybrids. Three major phenotypical hybrid forms were identified: GW-like, SW-like, and intermediate, with first two forms being more frequent in the central and south of Europe respectively. Whereas proportion of the intermediate form was relatively constant across the whole range of hybrid occurrence. The results indicate that hybrids of the two species have become significant part of woodpecker populations and thus should be considered in research and nature conservation projects.

## 1. INTRODUCTION

Woodpeckers in general are prone to hybridise due to the relatively similar morphological and behavioural characteristics of closely related species (Bakai et al. 2025a). The vast majority of woodpeckers inhabit wooded habitats and are dependent on trees for foraging and nesting (Winkler et al. 1995). Adaptations that allow them to excavate wood, together with a common ancestry (genetic similarities), determine their similar morphology (Short 1982, Prum et al. 2015). These similarities, along with sympatry or parapatry in many sister taxa, facilitate interspecific breeding. Recent systematic reviews have revealed that approximately one-fifth of the approximate 240 woodpecker species globally, are able to hybridise (Ottenburghs & Nicolaï 2024). That lights a new perspective on evolution of these birds and could have a consequences on ecology and reproductive biology of hybridising species. Unfortunately, until now, only a few genera are well examined in this respect, such as the North American *Sphyrapicus* and *Melanerpes* (Seneviratne et al. 2016, Barrowclough et al. 2017). Most of hybridising species are young in the evolutionary timescale, having evolved during the Pleistocene (Shakya et al. 2017). Interestingly, it seems that relatively more woodpecker taxa hybridise in the New World than in the Old World (Bakai et al. 2025a). Several genera are especially rich in hybridising species, such as the *Sphyrapicus*, *Melanerpes*, *Picumnus,* and *Dryobates* in the Americas, whereas in Eurasia, only some *Dendrocopos* regularly form interspecific pairs (Bakai et al., 2025a). In the latter genus, the taxa most prone to interbreeding are in the *D. major* complex: Sind (*D. assimilis*), White-winged (*D. leucopterus*), Great Spotted (*D. major*), and Syrian (*D. syriacus*) Woodpeckers (Gorman 2014). In addition, White-backed Woodpecker (*D. leucotos*) very occasionally hybridises with Great Spotted Woodpecker, too. GW and SW have a wide area of sympatric distribution due to the range expansion of SW which began in the 19^th^ century, when it began to move into Europe from its native range in Iran, Anatolia and the Levant (Michalczuk 2014). This species first settled in the Balkans and later expanded to the Pannonian Basin and Pontic region. Finally, SW established breeding populations in Austria, Czech Republic, Slovakia, Poland, Belarus and European Russia (Michalczuk 2014). A further expansion to the west and north was probably hindered by the unfavourable climate (wet winters), as it is more adapted to continental conditions (Keller et al. 2020). Recently, a decline in local SW populations, such as in Poland, have been observed (Michalczuk & Michalczuk 2015, Kajtoch & Kusal 2023).

When met, SW and GW express the various mutual interactions. Both exhibit some similar behavioural traits, particularly in aggressiveness when defending mates and territories. It seems that neither species is “stronger” or more aggressive than the other and dominance is a matter of context and individual bird “personality” (Figarski 2017). There are also cases, though rare, of these species showing cooperative behaviour when nesting; most likely caused by the loss of broods (Melnikov 2015). The nature of interaction between these co-occurring species is highly influenced by the availability of suitable nesting sites and food resources, two important determinants of woodpeckers distribution (Michalczuk & Michalczuk 2023). These factors differ considerably between the two main categories of synanthropic landscapes, namely rural and urban. Cases of polyandry between these two woodpeckers have been reported, however, extra-pair copulations are not well documented (Bakai et al. 2025b). The most interesting interaction between SW and GW is when they form interspecific pairs and hybrid young are produced (Gorman 1997). Such hybrids are fertile and an increasing number of backcrosses have been reported along with evidence of their subsequent successful breeding (Kajtoch & Kusal 2022). A recent summary of our knowledge on this phenomenon revealed that at least 3.6% of individuals in sympatric populations are hybrids, that number appears to be greater when dead birds are examined (Figarski & Kajtoch 2018). Indeed, thanks to the use of molecular markers it has been found that in some areas approximately 20% of birds in sympatric populations can be of interspecific origin (Michalczuk et al. 2014, Gurgul et al. 2019).

All of the above suggests that the hybridisation of these two woodpeckers is not uncommon. In the past, records of hybrids were mostly anecdotal and restricted to faunistic reports from the Balkans, Hungary and Slovakia (Winkler 1971). More recently, some of the best known examples are from Poland (Dudzik & Polakowski 2011). Nevertheless, extend of the European range of GW and SW hybrids remains unknown. It is unclear whether such hybrids are indeed present only along the edges of the SW’s range in Europe or are also present within the core area of its distribution (such as in the Balkans) and within its native range in the Near East (for example, in Turkey and Iran). A relatively untapped source of information are photographs available in online ornithological or biodiversity portals/databases, most of which have been collected via citizen science. Hence, we also took a chance to utilise this source.

Here, we present a summary of the current state of knowledge on the distribution of SW and GW hybrids and hybrid-like individuals. The main goal of the study was to show where hybrids had been detected so far. Additionally, we examined numerous images to assess whether all hybrids show similar phenotypic plumage features and whether there are different hybrid morphological forms. These data were then used to evaluate the hypothesis that hybrid-like individuals are not randomly distributed in Europe, but prevail along the edges of the range of SW, where the species is much less common and therefore a deficiency of mates forces birds to look for alternative partners. Secondly, we examined whether females are more frequent among hybrids after considering some previous findings from Poland where a high frequency of SW females was found among mixed pairs (Figarski & Kajtoch 2018). Thirdly, we investigate if the three phenotypic forms of hybrids (GW-like, SW-like and intermediate) are randomly distributed across Europe. Finally, the above listed goals were investigated in separately for populations inhabiting urban and rural landscapes, as differences in the structure of woodlands and availability of food resources between them can affect interactions between the two species and their hybrids.

## 2. METHODS

### 2.1 Citizen science

We assessed data from three widely-used bird databases in Europe: Ornitho (*ornitho.pl, ornitho.at, ornitho.de*), eBird (Sullivan et al. 2009) and iNaturalist (iNaturalist contributors, iNaturalist 2025), as well as national databases from the Czech Republic (*avif.birds.cz*), Slovakia (*birding.sk*), Hungary *(birding.hu*), Romania (*rombird.ro*), Ukraine (*ukrbin.com* and *uabirds.org*) and Russia (*russia.birding.day*). Unfortunately, we were unable to find country-level ornithological portals/databases for most of the Balkan countries and Asian ones, for example Turkey, where SW and GW are also sympatric. We did not investigate social media platforms such as Facebook, as they are unreliable in this context and, in addition, there is no straightforward way to extract precise data from the mass of uploaded information they host. We were aware of the bias that could result from differences in how particular portals are used in different countries, with often no precise information about the number and activity of users, especially in respect to the woodpeckers for which we sought data. All records were checked through the visual inspection of photographs available online, or provided by portal administrators.

Very few records in iNaturalist were assigned as SW×GW hybrids. Therefore, we decided to visually inspect around 5000 photographs there which were named as SW. Subsequently, we found some individuals that had hybrid plumage characteristics.

In eBird, we searched for hybrids using their search tool that facilitates the selection of such birds. We also examined all woodpeckers assigned as unidentified SW or GW, which had photographs and, once again, found hybrid individuals.

Only the Polish and Austrian Ornitho portals allowed direct access to records of hybrid individuals by using search tools. Additionally, the managers of both portals kindly gave us access to their overall databases in order to extract more information and photographs.

In addition, we were also able to extract hybrid records from the Slovakian *birding.sk* and Romanian *rombird.ro* sites.

We inspected all the SW image galleries in the Czech *avif.birds.cz*, Ukrainian *ukrbin.com* and *uabirds.org*, Russian *russia.birding* and Hungarian *birding.hu*, as there were no tools in these portals that facilitated searches for hybrids. Owing to the high numbers, we were unable to check all the images of GW in these databases.

Additionally, several photographic records of hybrids were shared to us directly by birdwatchers as a result of citizen science activities and within our own project “Mixed pairs and hybrids of Syrian and GWs” (at: sites.google.com/view/hybridization-woodpeckers/). Data from a review of GW × SW hybrids in Poland (Dudzik & Polakowski 2011) were also included.

Records from the same person about the same bird at the same time and place, repeated in more than one database, were removed. Such records were identified in the developed database via the sorting of records based on date, site and observer. Also, records of most likely the same birds from the same site observed over a short period of time were considered to be one, and the same was done for individuals which had plumage features that allowed their assignment as the same bird (from a given place frequently visited by observers). We could not find other ways of avoiding multiple records of the same individual although, in truth, such cases were extremely rare.

### 2.2 Field surveys

In the period 2023–2025, field surveys on 356 transects across Central and Southeastern Europe (including: Poland, Ukraine, Slovakia, Czech Republic, Austria, Germany, Hungary, Romania, Bulgaria, Greece), were conducted with a methodology adopted from the standard monitoring methods developed for SW surveys (Michalczuk & Michalczuk 2006a, Michalczuk & Michalczuk 2006b, Michalczuk et al. 2011). We were unable to conduct surveys in eastern Ukraine and Russia, therefore, this region was excluded from some analysis (share of GWs, SWs and their hybrids). The method used allows recording of both species, as well as their hybrids, since their reaction to playback is comparable (Figarski 2017). Our previous observations (by T. Figarski, G. Gorman, Ł. Kajtoch) also supported the view that the use of SW playback alone is sufficient for the detection of GW as well as the hybrids of these two species.

Bird survey was conducted along 2.4 km (2023) or 1.5 km (2024–2025) transects established along habitats suitable for SW (Michalczuk & Michalczuk 2016, Kajtoch & Figarski 2017), as GW can occupy the same habitats, but not the other way around (Figarski & Kajtoch 2018). The exact location for transects were chosen based on previous information about the presence of the two species and/or the data obtained from observers. Transects were surveyed from the 15th of March to 20th of April, which corresponds with the peak period of territorial activity (Michalczuk & Michalczuk 2006a,b). Through each transect, survey points were set at every 300 m at which playback of SW drumming and calls were used. All recorded birds were identified visually with binoculars, and the majority were photographed, whenever possible a series of photographs per individual was taken.

### 2.3 Site selection

The transect surveys were carried out in four regions in five locations per country. Exceptions were in Poland, Czech Republic and Slovakia, where we were able to collect data from more locations than elsewhere and more transects were done in large towns and their surroundings. In Ukraine, only three locations, and in Greece only one location, were surveyed. Overall, the distance between the nearest locations was around 80 km. At each location four transects were surveyed, two in urban areas and two in nearby rural areas. This was done to check on any possible differences between rural and urban landscapes. Exceptions were the cities of Brno, Bratislava, Warsaw and Kraków, where surveys were conducted on the large number of transects scattered across the city and surrounding areas. For statistical purposes, data from all localisations were adjusted to the same sample size.

### 2.4 Biogeographical variation of data

To analyse the geographic distribution of records, phenotypes and sexes, we grouped records according to their location into well-defined geographic regions of Europe:

1. “southern” - Balkans (European Turkey, Greece, Bulgaria).
2. “central” - Pannonian Basin and southern slopes of the Carpathians (Romania-Transylvania, Hungary, Slovakia, eastern Austria, eastern Croatia).
3. “eastern” - Pontic region (southern Ukraine, southern Russia, eastern Romania) and East European lowlands (northern and eastern of Ukraine, central Russia, Belarus) – this region was not considered for analyses involving data collected on transects.
4. “northern” - area north of the Carpathian and Sudetes Mountains (Poland, western Ukraine).
5. “western” - area between the Sudetes and the Alps (Czech Republic and northern Austria) and western Europe (Germany, Netherlands).

All records of hybrids were assigned to either rural or urban landscape types, based on their location.

We used our own data from field surveys across Europe (from Poland to Greece, and from Germany to Ukraine) to describe and visualise geographical variation in the co-occurrence of GW and SW, as well as mixed pairs and hybrids.

### 2.5 Hybrid determination

In this study, hybrid individuals were determined by both field observation and photographs based on their phenotypes.

We followed the set of plumage characters of hybrids described in Gorman (1999) and Bakai et al. (2025b) (Fig. S1). Ultimately, there is no typical plumage “type” for hybrids so, based on our field experience and the assessment of hundreds of photographs, we assigned birds to three basic forms, mostly focusing on the two most visible features: the extent of the post-auricular stripe and the amount of white in the tail, though other characteristics (see below) were also considered when visible:

1. intermediate phenotype – post-auricular stripe approx. 50% length of the corresponding stripe in GW, usually intermediate amount of red on the nape in males, similar extend of black and white on external rectrices, usually pink undertail, weak streaking on the flanks and belly.
2. GW-like phenotype – post-auricular stripe with very a narrow break (not connected to back of the head), a generally small red nape patch in males, black colour dominating on the external rectrices, red undertail, unmarked underparts.
3. SW-like phenotype – noticeably short post-auricular stripe (sometimes hardly distinguishable from the pattern in SW), a large red nape patch in males, white dominating on the external rectrices, pinkish undertail, streaked flanks and belly.

There are, however, no strict limits to these three types, therefore assigning individual birds is often somewhat subjective, especially when not all features are visible.

These features were mostly determined in adults, as the identification of juvenile hybrids individuals is challenging as differences are often less pronounced and overlap also exists.

### 2.6 Statistical analysis

The proportion of hybrids, as well as “pure” SW and GW individual was compared between rural and urban populations and between the geographical regions defined above. Since the data had a Gaussian distribution, a Dirichlet regression (DirichletReg package - Maier 2014) and a generally linear mixed model (glmmTMB - Brooks et al. 2017), the package in R was used to conduct the statistical analysis. Location, meaning two (or more) paired urban and rural count transects located in a specific single region, was used as a random factor to limit the influence of local conditions on the model results. To examine whether the share of the three defined hybrid phenotypes differed between rural and urban populations, as well as among the geographical regions, a Multinomial log-linear model was generated using the nnet package (Venables & Ripley 2002). To investigate whether the sex proportion of hybrid individuals differed between rural and urban populations, as well as among the regions, generalised linear models (considering the binomial distribution of dependent variables) were generated using a stats R package (R Core Team 2025). Additionally, generalised linear models (GLM) were generated using a lme4 package (Bates et al. 2015), and contrasts were checked with emmeans (Lenth 2025) for hybrid phenotypes and the sex proportion of hybrids. To calculate the share of hybrids versus “pure” woodpeckers, only data from field surveys was used as information from citizen science sources were deemed inadequate for such a analyses. For the remaining analyses (forms and sexes) all available data were used. Plots were prepared using a ggplot2 package (Wickham 2016). All analyses and data visualisations were made using R (v 4.5.1; R Core Team 2025) and RStudio (Posit team 2024). Maps showing the distribution of mixed pairs and hybrids across Europe (both from field and citizen science data), as well those showing the proportions of SW, GW and their hybrids, in both rural and urban landscapes, were prepared using QGIS (2025).

## 3. RESULTS

### 3.1 Records of mixed pairs and hybrids based on citizen science databases and transect surveys

#### iNaturalist

A total of 40 images of birds with hybrid phenotypes were found in *iNaturalist*. Most were from Russia (19), followed by Ukraine (8), Poland (4), Czech Republic (2), Austria (2), Greece (2), Bulgaria (1), Hungary (1) and Croatia (1) (Fig. 1, Tab. S1).

**Figure 1.**
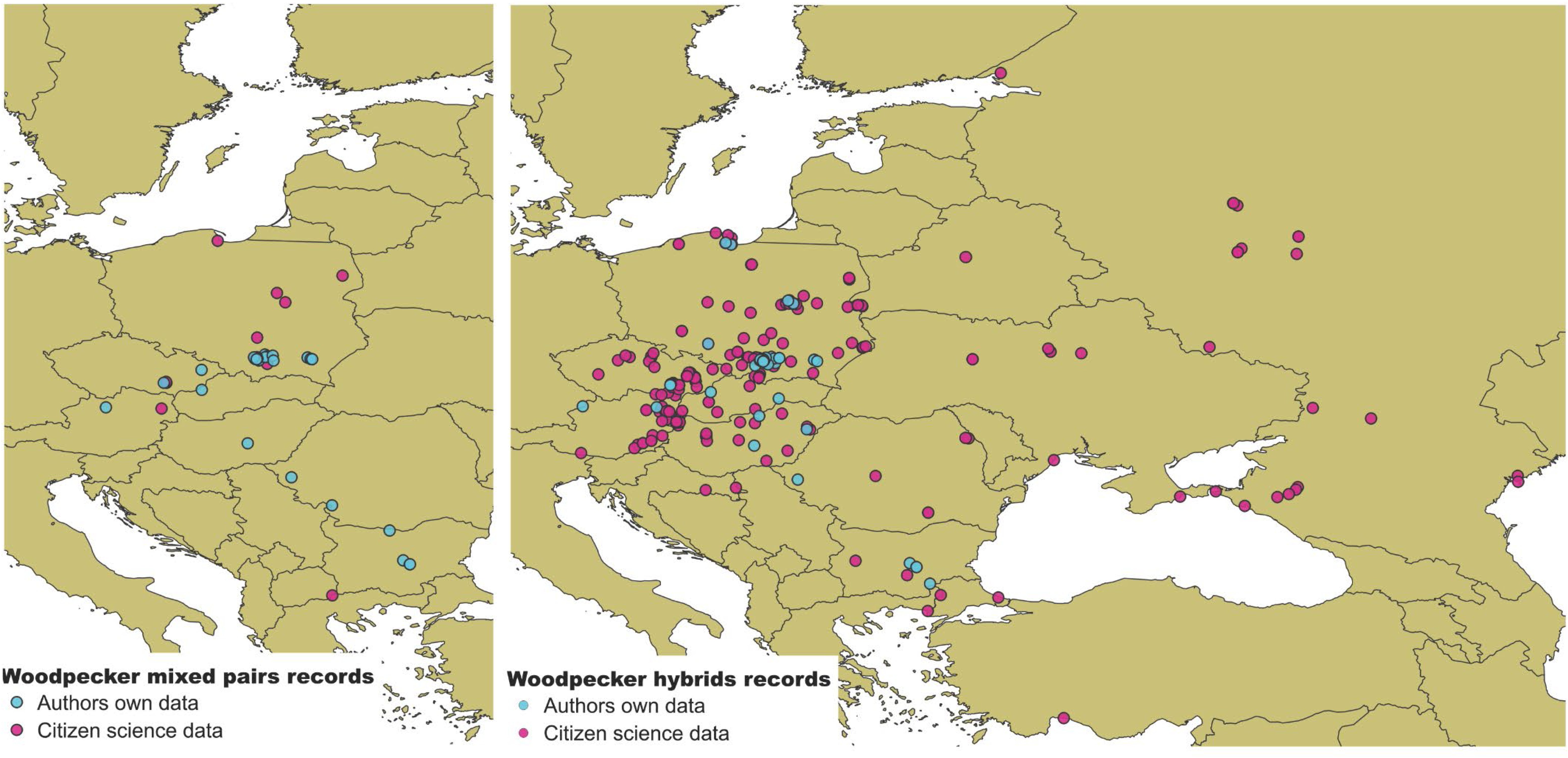
The records of mixed pairs (left) and hybrids (right) of SW and GW in Europe. Different colours indicate whether record originated from authors own data or citizen science.

One record of a mixed pair was found in Greece (Fig. 1, Tab. S2).

#### eBird

A total of 40 images of birds with hybrid phenotypes were found. Nearly half were from Poland (19), followed by Czech Republic (11) and Austria (3). Single records were also found for Belarus, Hungary, Croatia, Bulgaria, Greece, European Turkey and Anatolia (Fig. 1, Tab. S1).

Two records of mixed pairs were found in the Czech Republic (Fig. 1, Tab. S2).

#### Ornitho

A total of 84 of birds with hybrid phenotypes were found (excluding records that overlapped with data collected during our field surveys in Poland, see below): 54 from Poland and 30 from Austria (Fig. 1, Tab. S1).

One record of a mixed pair was found in Austrian Ornitho (Fig. 1, Tab. S2).

#### Avif.birds.cz

25 photographs of SW from the Czech Republic showed birds with hybrid phenotypes. Additionally, three reliable records without photographs were retrieved from this portal (Fig. 1, Tab. S1).

#### birding.sk

Hybrid phenotypes were found in six photographs from Slovakia (Fig. 1, Tab. S1).

#### birding.hu

Nine photographs of SW from Hungary showed birds with hybrid phenotypes (Fig. 1, Tab. S1).

#### rombird.ro

No records of hybrid-like woodpeckers were detected in this Romanian portal, although the number of available photographs was limited (Fig. 1, Tab. S1).

#### *ukrbin.com* and *uabirds.org*

Three photographs of birds with hybrid phenotypes were found in these two Ukrainian portals (Fig. 1, Tab. S1).

#### russia.birding

Hybrid-like individuals were found in nine photographs from Russia (Fig. 1, Tab. S1).

#### Other sources

There were additional 18 records from Poland (including 13 from the review of Dudzik & Polakowski (2011), 10 from Romania, two from the Czech Republic and one from Hungary provided by cooperating persons. Moreover, a photograph of a hybrid-like woodpecker in the Netherlands was discovered via internet search (Fig. 1, Tab. S1).

Six records of mixed pairs from Poland were found in Dudzik & Polakowski (2011) (Fig. 1, Tab. S2).

### Field survey data

A total of 54 hybrids were observed during field surveys across Europe during 2023-2025. Of these, 38 were in Poland, four in Bulgaria, three in the Czech Republic, three in Hungary, two in Slovakia, one in Romania, one in Austria and one in Germany (Fig. 1, Tab. S1).

Mixed pairs were also found in 31 sites across the transects, among them 20 in Poland, three in Bulgaria, two in Romania, two in the Czech Republic and one each in Slovakia, Hungary, Austria and Ukraine (Fig. 1, Tab. S2).

The list of all the records of hybrids and mixt pairs of SW and GW woodpeckers, retrieved from all citizen science databases and our own surveys, sorted by countries, is presented in Tab. S3.

### 3.2 Geographical distribution of hybrid forms

A total of 300 hybrid woodpeckers were recorded across all the investigated transects and in all citizen science databases. For 253 of these, we were able to determine the phenotype form, however, for nine records from citizen databases detailed information on the region was not available, hence the analysis was conducted on 244 birds.

Overall, most of the recorded hybrids had GW-like phenotypes (112 birds, 45.9%) followed by intermediate (71 birds, 29.1%) and SW-like (61 birds, 25%) phenotypes (Fig. 2, Fig. S2; Tab. S4). However, there was considerable variation in the phenotypic form distribution among the geographical regions.

**Figure 2.**
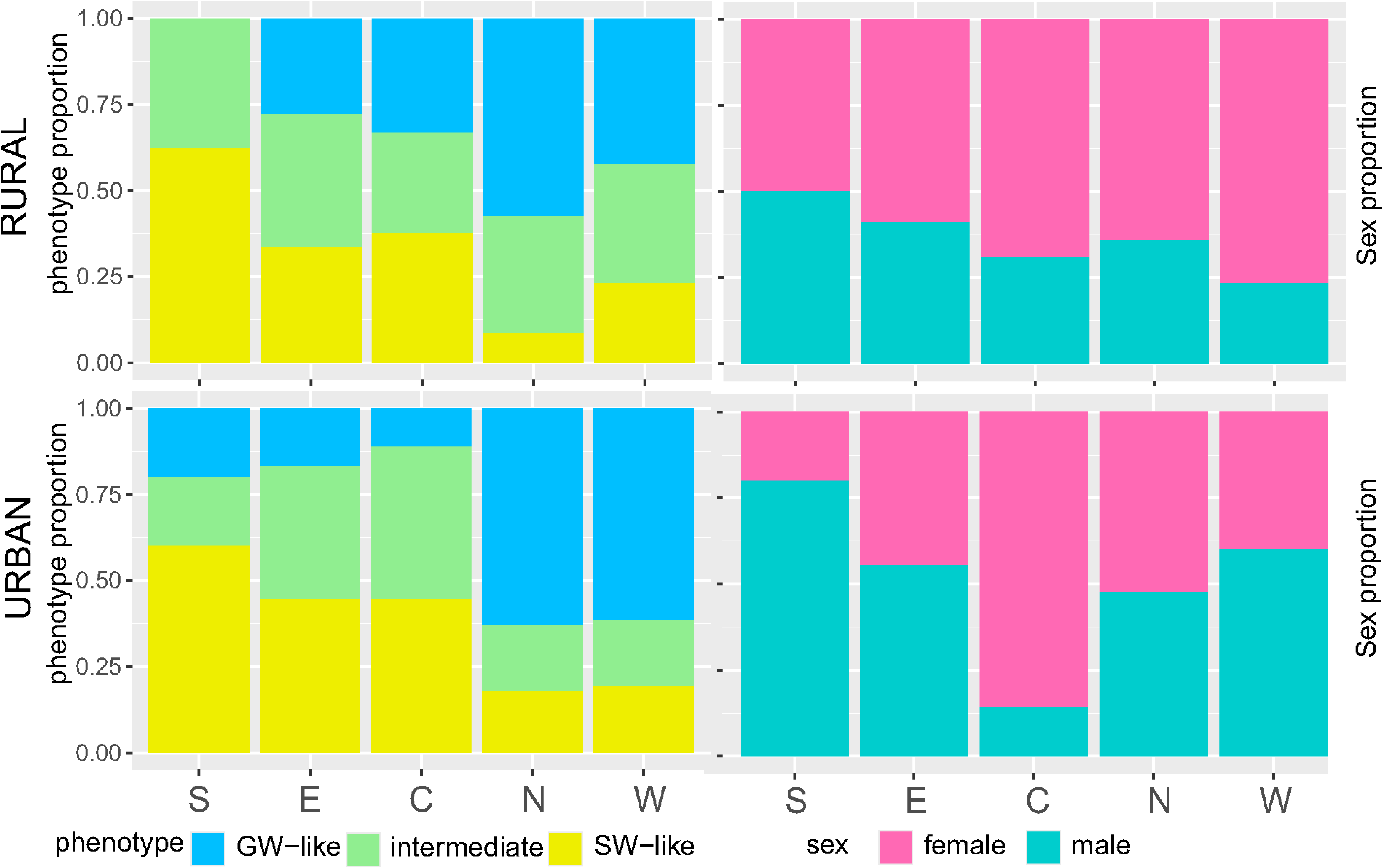
(left graph) Share between three hybrid phenotypes (GW-like, intermediate, SW-like) occurring in five regions (southern, central, eastern, western, northern) in rural and urban landscapes; (right graph) Sex ratio of hybrids across five regions (southern, central, eastern, western, northern) in rural and urban landscapes.

For instance, among 13 birds from the “southern” region 8 (61.5%) had SW-like phenotypes, 4 (30.8%) had intermediate phenotypes, and only one bird (7.7%) had the GW-like phenotype. In the “central” region, among 33 recorded hybrids the pattern of phenotypes was similar, though differences were less pronounced, with 13 (39.4%) birds having SW-like phenotypes, 11 (33.3%) intermediate and 9 (27.3%) GW-like phenotypes. Likewise, in the “eastern” region, out of 36 hybrids, SW-like with 14 (38.9%) and intermediate phenotypes also with 14 (38.9%) birds dominated, whereas the GW-like phenotype recorded in only 8 (22.2%) birds was the lest numerous. In contrast, in the “northern” (109 hybrids) and “western” (53 hybrids) regions the GW-like phenotype dominated, with 66 (60.5%) and 28 (52.8%) birds, respectively. Less common were the intermediate phenotypes 29 (26.4%) and 14 (26.4%), and SW-like phenotypes being the least numerous 14 (13.8%/12.8%) and 11 (20.8%), respectively.

Multinomial logistic regression confirmed that the three phenotypic forms of hybrids differ significantly in their relative share in respect to region (χ²=22.690, p=0.004), but also demonstrated no relationship to landscape (χ²=8.507, p=0.386) (Tab. 1).

**Table 1.** Summary of multinomial logistic regression models comparing the occurrence of Syrian Woodpecker and Great spotted Woodpecker hybrid phenotypic forms across regions and landscapes.

| models | $\chi^2$ | df | p value |
| --- | --- | --- | --- |
| Region | 22.69 | 8 | **0.004** |
| Landscape | 1.92 | 2 | 0.382 |
| Region:Landscape | 8.50 | 8 | 0.386 |
| „the best” Region | 38.35 | 8 | **<0.001** |

Moreover, generalised linear models (Tab. S5) indicated that GW-like phenotypes were significantly more frequent in northern and western regions, while SW-like phenotypes prevailed in southern, central and eastern regions. On the other hand, the frequency of intermediate phenotype hybrids did not differ within or across geographical regions, also the landscape type did not have a significant effect on the distribution on any of the phenotype forms.

### 3.3 Sex-ratio among hybrid woodpeckers

The overall sex ratio of hybrids was not equal as there was on average 1.4 females for each male (150 females : 107 males). However, this imbalance was apparently different across geographic regions. In “southern” populations there was 0.6 females for every male (N=13), while in the “central” region it was 2.7 females (N=33), in the “eastern” region 1.1 females (N=35), in the “northern” 1.4 females (N=121) and in the “western” 1.5 females per male (N=55) (Fig. 2, Fig. S2; Tab. S6). Yet, statistical analysis did not indicate a significant difference in sex ratio in respect to regions (Tab. 2, p = 0.503). But, it revealed a major difference in sex occurrence between landscapes (Tab. 2, p = 0.015). In urban landscapes the proportion was nearly equal with 0.97 females for each male (59 females : 61 males; N = 120) while in rural areas female hybrids were twice more abundant (91 females : 46 males; N = 137) .

**Table 2.** Summary of models comparing the proportion of male and female Syrian Woodpecker and Great spotted Woodpecker hybrids between regions and landscapes.

| models | df | Dev. | Resid.<br>df | Resid.<br>dev. | p value |
| --- | --- | --- | --- | --- | --- |
| Region | 4 | 3.337 | 5 | 8.743 | 0.503 |
| Landscape | 1 | 5.407 | 4 | 3.337 | **0.020** |
| Region:Landscape | 4 | 3.337 | 0 | 0.000 | 0.503 |
| „the best” Landscape | 1 | 5.924 | 8 | 6.157 | **0.015** |

### 3.4 Composition of birds in mixed pairs

A total of 48 mixed pairs of SW and GW was recorded both during our field survey (N = 31) and in the citizen science databases (N = 17). Data are summarized in Table S2.

GW (37.5%) tend to predominate within the mixed pairs compared to SW (31.5%) and hybrids (31.0%). The high frequency of GW occurrence is, however, clearly due to GW males which were found (over) 2.5 times more frequently (N=26, 27%) in mixed pairs than GW females (N=11, 10.5%). Compared to GW, males of SW (N=10, 10.5%) and hybrid woodpeckers (N=12, 13%) are considerably less frequent, in contrast to females of SW (N=20, 21%) and hybrid woodpeckers (N=17, 18%) which are found more frequently among the mixed pairs.

### 3.5 Co-occurrence of hybrids with their parental species

Our field observations in Europe indicate that in sympatric populations of SW and GW, SW numbers sharply decrease from S to N and from E to W (Fig. 3). GW shows opposite trends in its abundance (Fig. 3). SW is common mostly in the Balkans: Greece, Bulgaria and southern Romania, but they were also quite abundant in the Pannonian Basin in Hungary and S Slovakia. In NE Austria, Moravia, S Poland and Ukraine, SW still constituted 10–30% of local woodpecker populations, but farther to the N and W, GW were typically found alone, with exceptions of some localities in central Poland and Bohemia where SW were occasionally detected. The westernmost locality where SW was found was in E Germany in Görlitz. If a doubtful record in iNaturalist from Uckermark (NE Germany) from 2013 is excluded, this is only the third confirmed site with SW record in Germany, after Kronach in central Germany, 2016 and Frankfurt (Oder) in E Germany, 2022. Other western localities are in Prague (Czech Republic) and in Rottenhof near Linz (Austria).

**Figure 3.**
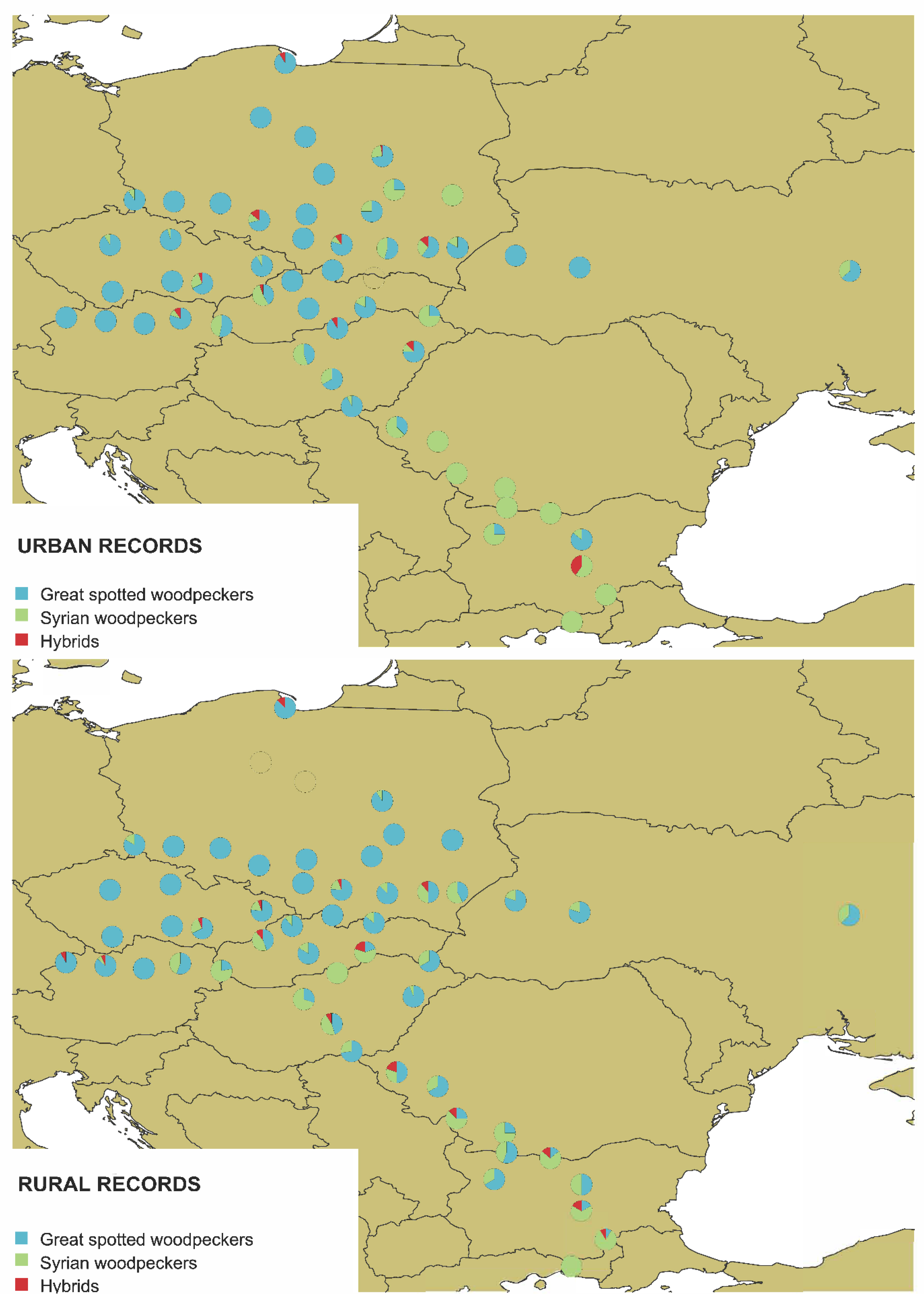
Co-occurrence of SW and GW and their hybrids in urban and rural populations, based on field surveys. Radial diagrams indicate the share of each species and hybrids in local populations.

Hybrid woodpeckers constitute a part of local GW and SW populations in several regions, for example: i) SE Poland, particularly in Kraków and Rzeszów and their environs, ii) Moravia, Czech Republic, and adjacent N Austria and W Slovakia, with a relatively higher number of hybrids found in Brno and its surroundings, iii) E Greece and E Bulgaria with the highest number found in Stara Zagora, and iv) SE Slovakia and E Hungary (Fig. 3). Locally, hybrids were also found in other cities such as Warsaw (central Poland), Opole (SW Poland) and Timisoara (NW Romania). The northernmost locality with hybrids was Gdańsk on the Polish Baltic coast. The westernmost locality was in Bad Fussing in SE Germany (the first confirmed record of hybrid in this country) (Fig. 3).

### 3.6 Frequency of GW, SW and their hybrids

Frequency of GW, SW and their hybrids was evaluated only based on data obtained from surveyed transects, data from citizen science databases were not included into that analysis.

Dirichlet regression revealed that the share of GW, SW and their hybrids differed significantly among regions (χ²=106.84, df=9, p<0.001), but not between landscapes (χ²=6.108, df=3, p=0.107), and that the effect of region and landscape combined was also insignificant ((χ²=14.187, df=9, p=0.116)).

Generalised linear mixed models were calculated for both species and hybrids to check how the proportion of given groups in their mixed populations differed among the four geographical regions (S/C/W/N) and two landscape types, considering interaction between two predictors (Tab. 3, Fig. 4). The model revealed a significant difference in GW and SW proportions for different regions but not for landscape types. The only significant difference between rural and urban landscapes was for SW in “southern” populations. There was no significant difference in hybrid proportions between regions or landscapes (Tab. 3). Although, some cities seemed to host high number of hybrids e.g. Kraków and Rzeszów in Poland, Brno in the Czech Republic, Debrecen in Hungary and Stara Zagora in Bulgaria. In rural landscapes, hybrids were present, for example, in S Poland, the Czech Slovak border, S Slovakia, E Romania and C Bulgaria (Fig. 3, Fig. 4).

**Figure 4.**
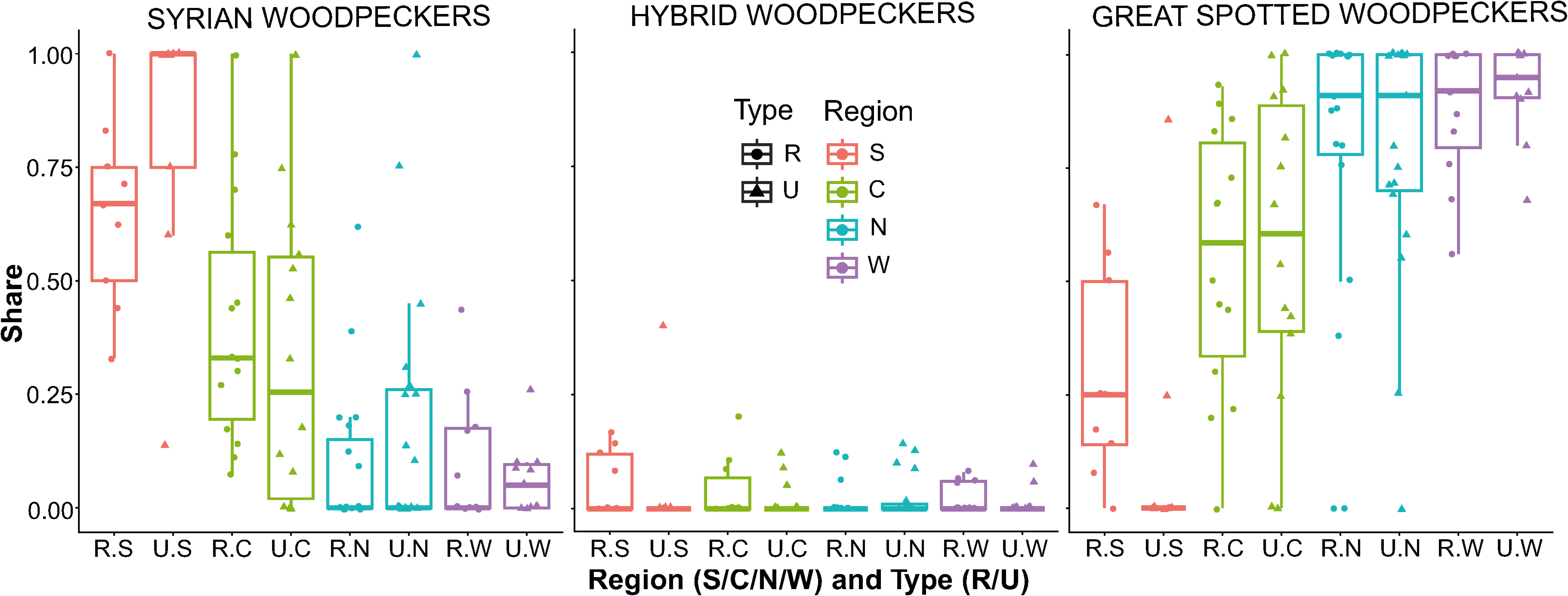
The boxplots show the proportion of SW, SW×GW hybrids and GW in four defined regions (S – southern, C – central, N – northern, W – western) and landscape type (R – rural, U - urban).

**Table 3.** Summary of generalised linear mixed model (GLMM) assessing the influence of region (S – southern, C – central, N – northern, W – western) and landscape type (R – rural, U – urban) as well as their interactions on Great spotted Woodpecker, Syrian Woodpecker and their hybrid proportions in surveyed populations. Probability values of significant factors (p≤0.05) are in bold.

| source of variation | estimate | std. error | z value | p value |
| --- | --- | --- | --- | --- |
| Great spotted Woodpecker |  |  |  |  |
| (Intercept) | 0.550 | 0.071 | 7.771 | **<0.001** |
| RegionN | 0.234 | 0.093 | 2.516 | **0.012** |
| RegionS | -0.259 | 0.113 | -2.291 | **0.022** |
| RegionW | 0.324 | 0.107 | 3.043 | **0.002** |
| LandscapeU | 0.028 | 0.100 | 0.285 | 0.776 |
| RegionN:LandscapeU | -0.014 | 0.131 | -0.103 | 0.918 |
| RegionS:LandscapeU | -0.196 | 0.160 | -1.226 | 0.220 |
| RegionW:LandscapeU | 0.021 | 0.151 | 0.139 | 0.889 |
| Syrian Woodpecker |  |  |  |  |
| (Intercept) | 0.407 | 0.061 | 6.627 | **<0.001** |
| RegionN | -0.314 | 0.081 | -3.876 | **<0.001** |
| RegionS | 0.245 | 0.098 | 2.488 | **0.013** |
| RegionW | -0.306 | 0.093 | -3.305 | **0.001** |
| LandscapeU | -0.076 | 0.079 | -0.963 | 0.335 |
| RegionN:LandscapeU | 0.159 | 0.104 | 1.533 | 0.125 |
| RegionS:LandscapeU | 0.257 | 0.126 | 2.033 | **0.042** |
| RegionW:LandscapeU | 0.037 | 0.119 | 0.311 | 0.756 |
| Hybrid |  |  |  |  |
| (Intercept) | 0.043 | 0.016 | 2.718 | **0.007** |
| RegionN | -0.028 | 0.021 | -1.336 | 0.182 |
| RegionS | 0.015 | 0.025 | 0.575 | 0.565 |
| RegionW | -0.018 | 0.024 | -0.761 | 0.447 |
| LandscapeU | -0.024 | 0.018 | -1.302 | 0.193 |
| RegionN:LandscapeU | 0.033 | 0.024 | 1.389 | 0.165 |
| RegionS:LandscapeU | 0.011 | 0.029 | 0.367 | 0.714 |
| RegionW:LandscapeU | 0.013 | 0.028 | 0.486 | 0.627 |

## 4. DISCUSSION

Here we show that both citizen science data and detailed field surveys with photographic documentation consistently help to confirm wide distribution of mixed pairs and SW×GW hybrids.

Previous knowledge on SW × GW hybrids was limited to local observations that suggested incidental cases of this phenomenon (Winkler 1971, Dudzik & Polakowski 2011). Although hybrids were quite abundant in some areas, recent studies based on field surveys and genetic research revealed that hybrids were particularly abundant in sympatric urban populations. For example, in southern Poland approximately 20% of SW and GW populations may be of interspecific origin (Figarski & Kajtoch 2018, Gurgul et al. 2019). Our data indicates that hybrids are rare but widespread, and their proportion compared to “pure” birds in the whole study area is ca. 3%. However, in some local populations where surveying has been regular (for example, Kraków and its surroundings) that number increases to 10%, and thus hybrids locally comprise a significant part of the woodpecker community.

Hybrids were detected across the whole European range of SW. Elsewhere, only one record from S Anatolia from 1999 (eBird) was found and, although there is no photographic evidence, the description of this museum specimen supports its hybrid origin. However, the lack of hybrid records from the Middle and Near East may be due to the sparsity of birdwatchers and hence citizen science efforts in the region. Another reason could involve the zoogeography of these two woodpeckers in that region, where these birds are mostly allopatric, occupying different habitats, with distribution mostly related to altitude: SW is common in lowlands, whereas GW occupies only mountain forests (Keller et al. 2020)). A consequence of that partial altitudinal segregation is that contact, and hence the opportunity to form mixed pairs, is limited.

Some regions in Europe have higher numbers of hybrid records than others. It is unclear if such “hot spots” are indeed centres of hybridisation or simply areas with more observers. However, some of these “hot spots” have been confirmed by both citizen science reports and by our field surveys. Hybrids have been found mainly on the edge of SW range in Poland, Czech Republic and Austria, as well as in SE Ukraine and S Russia. Another region with relatively higher numbers of hybrids was found in the Pannonian Basin (S Slovakia and E Hungary). Notably, most hybrid records in these “hot spots” were recorded along foothills and valleys close to mountains, in the Carpathians, Alps, Crimea and the Caucasus. In such regions, SW breeds in lowland rural and urban landscapes, whereas GW breeds mostly in montane forests with a tendency to spread to lower altitudes (Kajtoch & Figarski 2017). All this creates a transitional zone where both species can easily meet and potentially interbreed. Another “hot spot” is found in the border area of Bulgaria, Greece and Turkey, gateway from Asia to Europe where expansion of SW range into Europe was firstly recorded more than a century ago (Michalczuk 2014). Interestingly, hybrids have been found in both urban and rural populations in Europe, although the share of hybrids differs across the range of SW. It seems that some urban populations form local “hot spots” where interspecific mating accelerates, but in some rural areas, too, hybrids can frequently occur. It is likely that the reason for this is related to both the availability of mates and the distribution of suitable habitats for both species (e.g. parks in cities, orchards in rural areas for SW and (urban or rural forests and woodlands for GW). Forest populations of GW (where SW tends to be absent (Lešo et al. 2019) is usually free of hybrids, although a few records were recently found in forests around Kraków in S Poland (Kajtoch & Kusal 2023, Kajtoch et al. 2025). Therefore, forests should not be omitted in search for hybrids.

The first records of SW×GW hybrids in Europe hail from the Balkans and the Pannonian Basin, about 50 years after SW occupied Europe (Winkler 1971, Gorman 1997). Nevertheless, it is likely that hybridisation began in parallel with SW expansion already the end of the 19^th^ century, but it was unnoticed. If that assumption is correct, the gene flow between GW and SW may well have a longer and more significant impact than often thought. Apart from the evolutionary and ecological consequences of this phenomenon, there are also other effects that have usually been neglected by ornithologists.

Firstly, most scientific research on woodpeckers in Europe has not considered the presence of hybrids, and such individuals were usually omitted or intentionally ignored during field surveys. Perhaps, it is easier for researchers to avoid such individuals, as their presence complicate interpretation of the results. By considering hybrids together with “pure” birds, rather than separately, researchers do not account for the different genetic and phenotypic characters of hybrids that likely make them and “pure” individuals respond differently to environmental factors and possibly also to the presence of each other. Having alleles from two genomic pulls, hybrids are characterized by higher heterozygosity, which might give them advantages such as broader habitat tolerance. This increased heterozygosity may also result in a hybrid superiority that allows such birds to outcompete their parental species. It seems that in case of SW×GW hybrids, they might replace the SW in most habitats (Kajtoch & Kusal 2023).

In some populations a deficiency of SW individuals accelerates hybridisation. Such deficiencies mostly concern males, as previous studies have proved that mixed pairs are constituted mainly by SW or hybrid females mating with GW males (Figarski & Kajtoch 2018). This could also explain the higher share of females over males among hybrids. But we also cannot exclude, that hybrid females are more attractive partners, however, potential mechanisms underlying such preferences are unknown.

Additionally, neglecting hybrids can be a significant problem when monitoring woodpeckers, particularly in case of SW in its peripheral populations where it is rare. SW is annexed in the Bird Directive of European Union, and special protection areas for their conservation are designated (Directive 2009/147/EC). Lack of information about the number of hybrids makes its protection less effective. Unfortunately, European Union legislation does not consider hybrids at all, so their status is unclear. In addition, bird monitoring programs in the EU do not take into account application of playback, which was proved in our research to be a very effective tool in detection of breeding SW. Thus, the current methodology, unfortunately does not provide the best tools for monitoring and protection of that species.

Another interesting finding is that apart from the most apparent hybrid individuals, that have distinguishable intermediate characters between SW and GW (often the first generation hybrids), we have also documented existence of two other types of hybrids. One mainly resembles GW (GW-like), whereas other has more characters of SW (SW-like). It is likely that differences in plumage reflect the contribution of alleles of their parental species and continue in further generations to produce “backcrosses”. Support for this hypothesis lies in the distribution of these two types of hybrids. Individuals with a majority of GW characters were mostly found in the outermost localities of the range of SW, from Moravia, through Poland to central Russia. In contrast, individuals with a dominance of SW plumage features were found mostly in the Balkans, Romania and the Pontic region (Fig. 3). While birds with a variable share of features from parental species were found across the whole investigated area, the dominance of either SW or GW plumage seems to be highly related with the regional abundance of each species. This seems to be a clear evidence that genome contribution is responsible for plumage patterns but further genetic studies across the whole distribution of SW×GW hybrids is needed to confirm this hypothesis. GW-like hybrids dominated over birds with SW-like and intermediate plumages, although all of them were present in woodpecker populations (Table S4). This indicates that there is no single dominating type, which means that the mating pattern of hybrids (especially backcrosses) is determined by the availability of partners, whether pure SW, pure GW or hybrids. This situation fits the so-called “hybrid swarm” population structure (Fitzpatrick & Shaffer 2007). Such swarms are most likely present in south-eastern Poland and Moravia, but a concentration of records in other areas, such as Burgenland in E Austria, the Slovakia-Hungary border, around the Azov Sea and in the SE Balkans, suggest existence of additional “hot spots” that deserve study. The presence of hybrids should be considered during ornithological research and monitoring programs conducted in these areas.

Frequent presence of hybrids in some populations leads to the question of how to find and identify them. There are three reliable ways for identification of hybrids. The first is to find a mixed pair of SW and GW and document characteristics of their offspring. However, such strategy requires extensive effort and time, making it not effective method for large-scale surveys or monitoring. The second is to apply molecular markers, as it has been proven that mitochondrial DNA, nuclear introns, microsatellites and genotyping based on numerous single nucleotide polymorphism, are all accurate in detecting both first-generation hybrids and backcrosses (Michalczuk et al 2014, Gurgul et al. 2019). But this method, too, is also ineffective for large-scale surveys and monitoring due to the need to collect biological samples such as feathers or blood. Therefore, though less effective, but still reliable is visual identification, with detailed phenotype examination, specifically differences in plumage. A set of plumage features that allows the distinct identification of hybrid SW×GW has been published (Gorman 1999, Bakai et al. 2025b). Usage of a high-quality digital camera, and collection of series of photographs is particularly effective and provides good quality documentation for further evaluations. However, some individuals with minor plumage characteristics specific to hybrids still cannot always be safely assigned as hybrids.

Furthermore, aberrant individuals cannot always be ruled out. Then again, all of the aberrant pied woodpeckers reported to date in the literature have been either melanistic or leucistic – full or partial (Karlík & Veľký 2007, Olszewski 2007, van Grouw et al. 2021). Therefore, a bird with intermediate or mixed plumage features is more likely to be of mixed ancestry with its “unusual” plumage caused by gene introgression. Another problem is that relatively few birdwatchers are aware of the existence of woodpecker hybrids and therefore they can be easily overlooked, as it is often the case even during professional ornithological research (see Bakai et al. 2025b). The increasing availability of high-quality optical devices has probably aided the documentation of hybrids. Furthermore, the growth of citizen science has helped in determining hybrids as today many birdwatchers carry high-quality cameras and share their images on online portals such as iNaturalist, eBird and Ornitho. These databases provide invaluable sources of data, although all records need to be verified according to current scientific knowledge and misidentified bird records corrected (Sullivan et al. 2009). Still, the thousands of photographs available in both international and country-level portals and websites are good sources of data that allow us to extend our knowledge on occurrence and geographical distribution of SW×GW hybrids.

## 5. Conclusions

Our research supports the hypothesis that SW×GW hybrids are more widespread across Europe than previously thought. The majority of records are from the edges of SW range in central and eastern Europe, but not exclusively: local hybrid “hot spots” seems to exist also in other regions, such as Balkans, Carpathian Basin and the Pontic region. Single cases of hybrids have also been found further to the W and N outside of the SW breeding range. Moreover, hybrids are equally widespread in urban and rural landscapes, although there are regional differences.

The morphology (plumage patterns and colours) of hybrids varies and the mixture of features is likely determined by contribution of genomes from both SW and GW. SW-like hybrids dominate in the core part of the European range of SW (i.e., Balkans, Pannonian Basin, Pontic region) but GW-like hybrids are more common along the edges of the SW range. The various forms of hybrids that are found across Europe suggest that hybridisation is a continuous process and that backcrosses are able to breed and are fertile.

Females dominate among SW and GW hybrids, but the underlying mechanism, whether this pattern is driven by genetic, behavioural or fitness factors, remain unclear. A possible explanation may reside in the different phenotypes of hybrid males and females, with males resembling more the parental species and thus being hard to identify. Further studies are needed to understand this population trend.

Finally, it should be pointed that most of the above-mentioned hybrid records were assigned as such using field identification. However, the most reliable way, and possibly the only certain way, to identify hybrids is to analyse the genetics of the birds in question. This was the method during some previous investigations (Michalczuk et al. 2014, Gurgul et al. 2019, Kajtoch & Kusal 2023). A project that aims to shed more light on the topic of SW and GW hybridisation is ongoing. Our work demonstrates that the use of citizen science can also provide significant data that are complimentary to research on SW and GW hybridization.

## Acknowledgements

This paper was prepared as a part of a grant funded by the National Science Centre, Poland (project number UMO-2022/47/O/NZ9/02044, granted to Ł. Kajtoch).

We are grateful to the following for their contributions to field surveys and data sharing: Krzysztof Basista, Johanka Bláhová, Jan Čapek, Peter Chrašč, Balázs Csibrány, Krzysztof Czarnocki, Adam Dmoch, Ryszard Dworak, Paweł Grochowski, Fatima Hayatli, Lucie Hornátová, Mariusz Godlewski, Marta Gołek, Tomasz Janiszewski, Bartek Łukaszewicz, Katarzyna Kusal, Maria Madej, Konrad Malec, Magdalena Naber, Samuel Pačenovský, Bartłomiej Stankiewicz, Daniela Svojanovská, Patryk Rowiński, Jan Špička, Zbyszek Swiacki, Marta Świtala, Krzysztof Tabernacki, Matěj Tvarůžka, Łukasz Wardecki, Marta Wołoszyn, Jakub Wyka, Lukas Vana, Tomasz Ziółkowski, Paweł Żarkiewicz, Karolina Żukowska,

Moreover, we are thankful for all the databases for making bird records publicly available: ornitho.pl, ornitho.at, ornitho.de, inaturalist.org, avif.birds.cz, birding.sk, birding.hu, rombird.ro, ukrbin.com, uabirds.org and russia.birding. Special thanks are due to the manager of the Austrian and the Polish ornitho portals for allowing direct access to all records with all details (photographs, localisations, etc.).

---

**Fig. S1.**
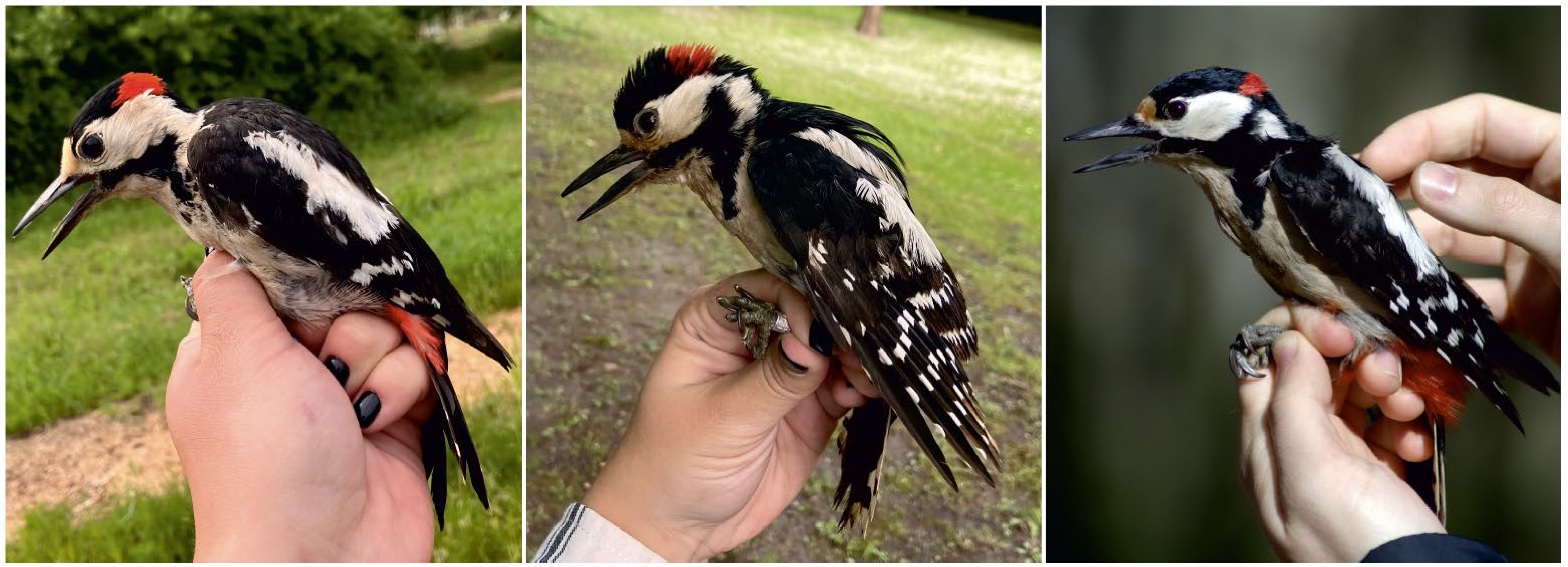
Photographs of Syrian woodpecker (left), hybrid (middle) ànd Great-spotted woodpecker (right) (photo: Zuzanna Pochłopień).

**Figure S2.**
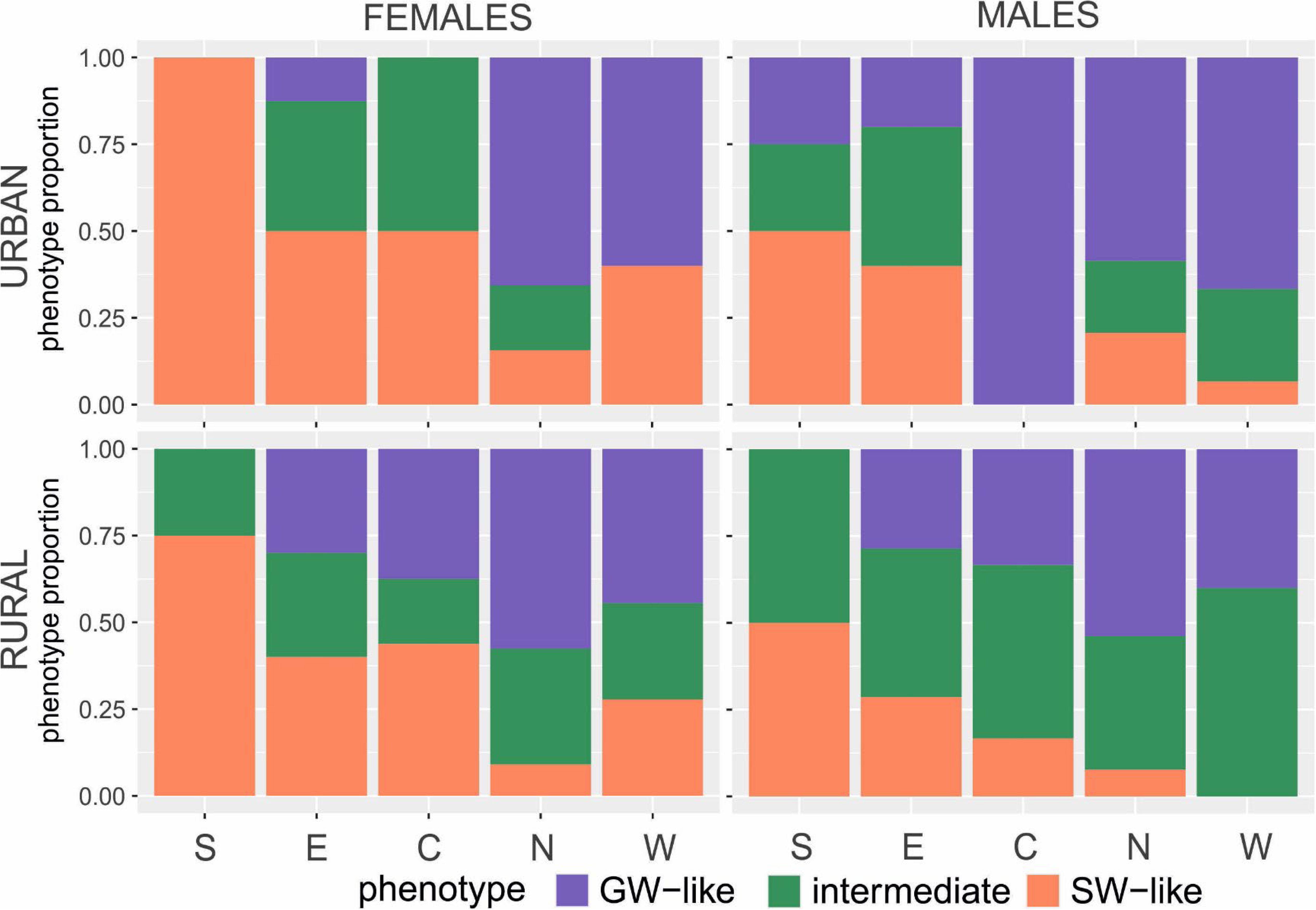
Proportion of three forms of hybrid woodpeckers (Great spotted – like, GW-like; Syrian – like, SW-like; intermediate), and sexes in hybrid woodpeckers across regions: southern (S), eastern (E), central (C), Northern (N) and western (W) and landscapes: urban (U) and rural (R).

**Table S1.**
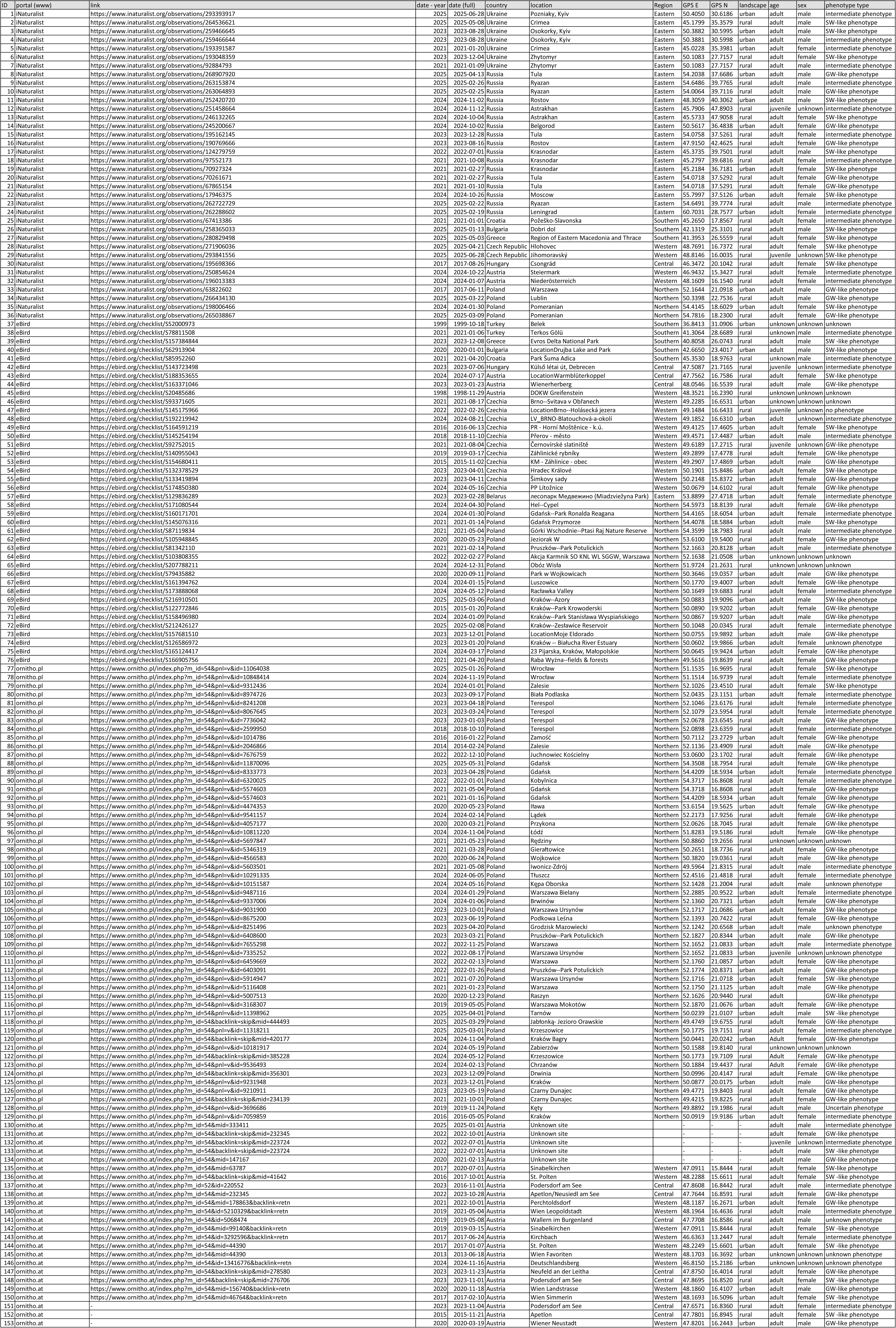
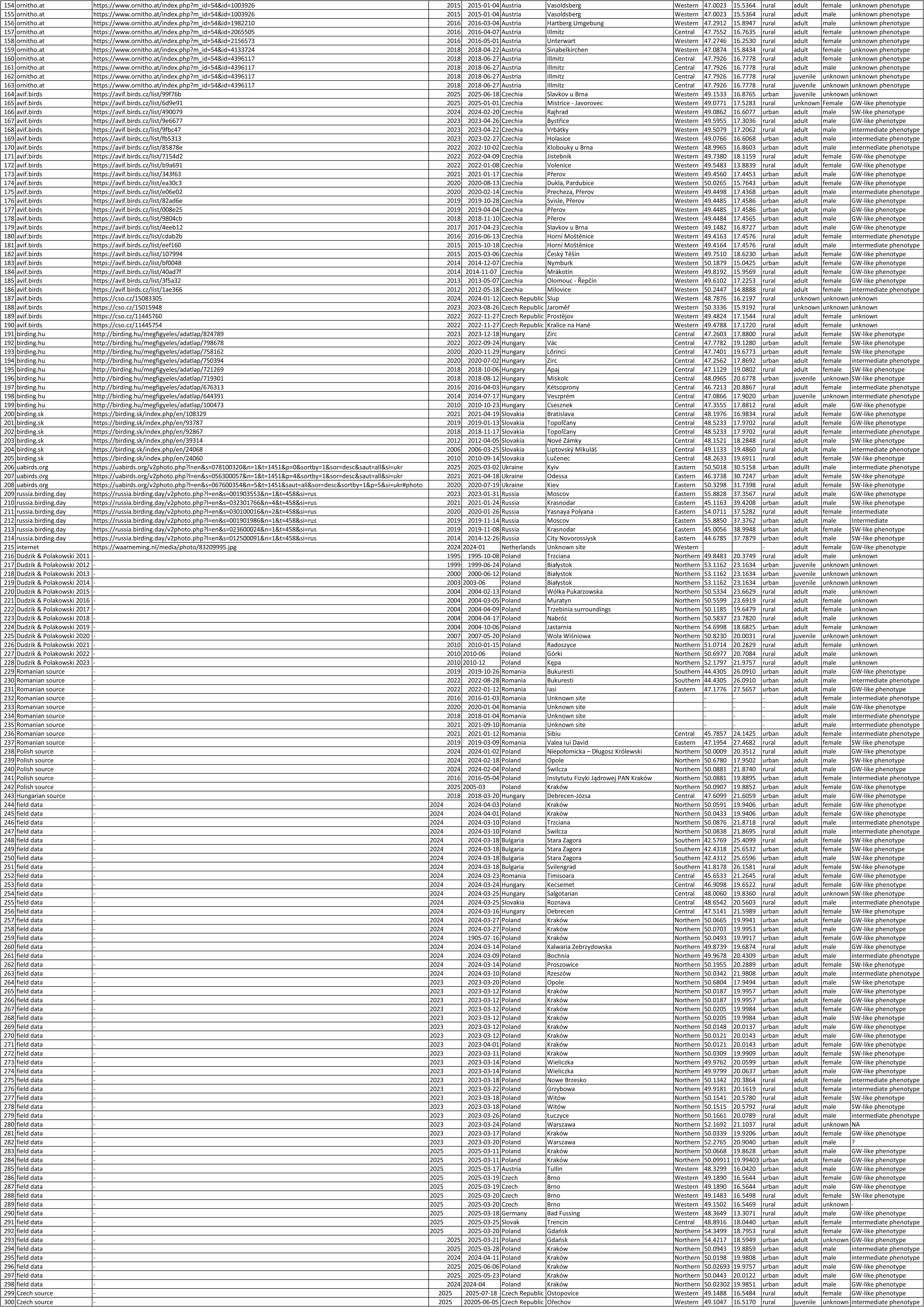
Records of hybrids.

**Table S2.** Records of mixed pairs.

| ID | portal (www) | link | date - year | date (full) | country | location | GPS E | GPS N | GW | SW | hybrid |
| --- | --- | --- | --- | --- | --- | --- | --- | --- | --- | --- | --- |
| 1 | ornitho.at | https://www.ornitho.at/index.php?m_id=548 | 2019 | 2019-05-08 | Austria | Wien | 48.1964 | 16.4636 | female | male |  |
| 2 | eBird | https://ebird.org/checklist/S145175966 | 2022 | 2022-02-26 | Czech R. | Lukáš Váňa | 49.1484 | 16.6433 | male | female |  |
| 3 | eBird | https://ebird.org/checklist/S192219942 | 2024 | 2024-08-21 | Czech R. | Lukáš Váňa | 49.1852 | 16.6309 | male | female |  |
| 4 | iNaturalist | https://www.inaturalist.org/observations/22 | 2024 | 2024-06-06 | Greece | unknown | 41.2795 | 22.7846 |  | male | female |
| 5 | Dudzik & Polakowski 2011 | - | 1980 | 1980-06-01 | Poland | Trzciana | 49.8483 | 20.3749 | male | female |  |
| 6 | Dudzik & Polakowski 2011 | - | 1998 | 1998-03-26 | Poland | Warszawa | 52.1287 | 21.0458 | male | female |  |
| 7 | Dudzik & Polakowski 2011 | - | 2001 | 2001 | Poland | Gdańsk | 54.4054 | 18.5421 | female | male |  |
| 8 | Dudzik & Polakowski 2011 | - | 2003 | 2003-06-07 | Poland | Białystok | 53.1162 | 23.1633 | male | female |  |
| 9 | Dudzik & Polakowski 2011 | - | 2004 | 2004-03-13 | Poland | Pomiechówek | 52.4750 | 20.7350 | male | female |  |
| 10 | Dudzik & Polakowski 2011 | - | 2007 | 2007-05-20 | Poland | Wola Wiśniowa | 50.8229 | 20.0031 | male | female |  |
| 11 | Kajtoch & Kusal 2023 | - | 2014 | 2014-04 | Poland | Kraków | 50.0123 | 20.0064 | male | female |  |
| 12 | Kajtoch & Kusal 2023 | - | 2015 | 2015-04 | Poland | Wieliczka | 49.9819 | 20.0632 | female |  | male |
| 13 | Kajtoch & Kusal 2023 | - | 2021 | 2021-04 | Poland | Kraków | 50.0908 | 19.9699 | male | female |  |
| 14 | Kajtoch & Kusal 2023 | - | 2021 | 2021-04 | Poland | Kraków | 50.0195 | 19.9975 |  |  | male, female |
| 15 | Kajtoch & Kusal 2023 | - | 2021 | 2021-04 | Poland | Wieliczka | 50.0021 | 20.0838 | male | female |  |
| 16 | Kajtoch & Kusal 2023 | - | 2021 | 2021-04 | Poland | Wieliczka | 49.9799 | 20.0629 |  | male | female |
| 17 | Kajtoch & Kusal 2023 | - | 2022 | 2022-04 | Poland | Kraków | 50.0168 | 19.9949 |  |  | male, female |
| 18 | field data | - | 2024 | 2024-03-10 | Poland | Swilcza | 50.0876 | 21.8718 | female |  | male |
| 19 | field data | - | 2024 | 2024-03-10 | Poland | Swilcza | 50.0838 | 21.8695 |  | female | male |
| 20 | field data | - | 2024 | 2024-03-17 | Bulgaria | Nikopol | 43.6812 | 24.9061 | male | female |  |
| 21 | field data | - | 2024 | 2024-03-18 | Bulgaria | Kazanlyk | 42.5769 | 25.4099 |  | male | female |
| 22 | field data | - | 2024 | 2024-03-18 | Bulgaria | Stara Zagora | 42.4318 | 25.6532 |  | male | female |
| 23 | field data | - | 2024 | 2024-03-18 | Bulgaria | Stara Zagora | 42.4312 | 25.6596 |  | female | male |
| 24 | field data | - | 2024 | 2024-03-22 | Romania | Poroina | 44.6137 | 22.7744 | female | male |  |
| 25 | field data | - | 2024 | 2024-03-23 | Romania | Timisoara | 45.6519 | 21.2727 | male | female |  |
| 26 | field data | - | 2024 | 2024-03-24 | Hungary | Kecsemet | 46.9098 | 19.6522 | male |  | female |
| 27 | field data | - | 2024 | 2024-03-27 | Poland | Kraków | 50.0665 | 19.9941 | male |  | female |
| 28 | field data | - | 2024 | 2024-03-27 | Poland | Kraków | 50.0731 | 19.9953 | female |  | male |
| 29 | field data | - | 2024 | 2024-04-11 | Poland | Kraków | 50.0493 | 19.9917 | male |  | female |
| 30 | field data | - | 2024 | 2024-03-14 | Poland | Proszowice | 50.1955 | 20.2889 |  | male | female |
| 31 | field data | - | 2024 | 2024-03-10 | Poland | Rzeszów | 50.0342 | 21.9808 | female |  | male |
| 32 | field data | - | 2024 | 2024-03-10 | Poland | Rzeszów | 50.0238 | 22.0401 | male | female |  |
| 33 | field data | - | 2023 | 2023-03-12 | Poland | Kraków | 50.0121 | 20.0143 | male |  | female |
| 34 | field data | - | 2023 | 2023-03-14 | Poland | Wieliczka | 49.9799 | 20.0637 | male |  | female |
| 35 | field data | - | 2023 | 2023-03-18 | Poland | Nowe Brzesko | 50.1342 | 20.3864 | male |  | female |
| 36 | field data | - | 2023 | 2023-03-19 | Poland | Wawrzeńczyce | 50.1010 | 20.3049 | male | female |  |
| 37 | field data | - | 2023 | 2023-03-18 | Poland | Witów | 50.1541 | 20.5780 |  | male | female |
| 38 | field data | - | 2023 | 2023-03-17 | Poland | Kraków | 50.0339 | 19.9206 | male |  | female |
| 39 | field data | - | 2023 | 2025-03-11 | Poland | Kraków | 50.0991 | 19.9940 | male |  | female |
| 40 | field data | - | 2025 | 2025-03-16 | Poland | Brzesko | 49.9721 | 20.6069 | male | female |  |
| 41 | field data | - | 2025 | 2025-03-18 | Austria | Linz | 48.2408 | 14.4040 | male | female |  |
| 42 | field data | - | 2025 | 2025-03-20 | Czech | Brno | 49.1502 | 16.5469 | male |  | female |
| 43 | field data | - | 2025 | 2025-03-20 | Czech | Jicin | 49.6284 | 17.9445 | female | male |  |
| 44 | field data | - | 2025 | 2025-03-25 | Slovak | Trencin | 48.8888 | 17.9520 | male | female |  |
| 45 | field data | - | 2025 | 2025-05-23 | Poland | Kraków | 50.0443 | 20.0122 | female |  | male |
| 46 | field data | - | 2024 | 2024-04-11 | Poland | Kraków | 50.0198 | 19.9808 | female |  | male |
| 47 | field data | - | 2025 | 2025-03-28 | Poland | Kraków | 50.0943 | 19.8859 | female |  | male |
| 48 | field data | - | 2024 | 2024-04 | Poland | Kraków | 50.0230 | 19.9851 |  | female | male |

**Table S3.**
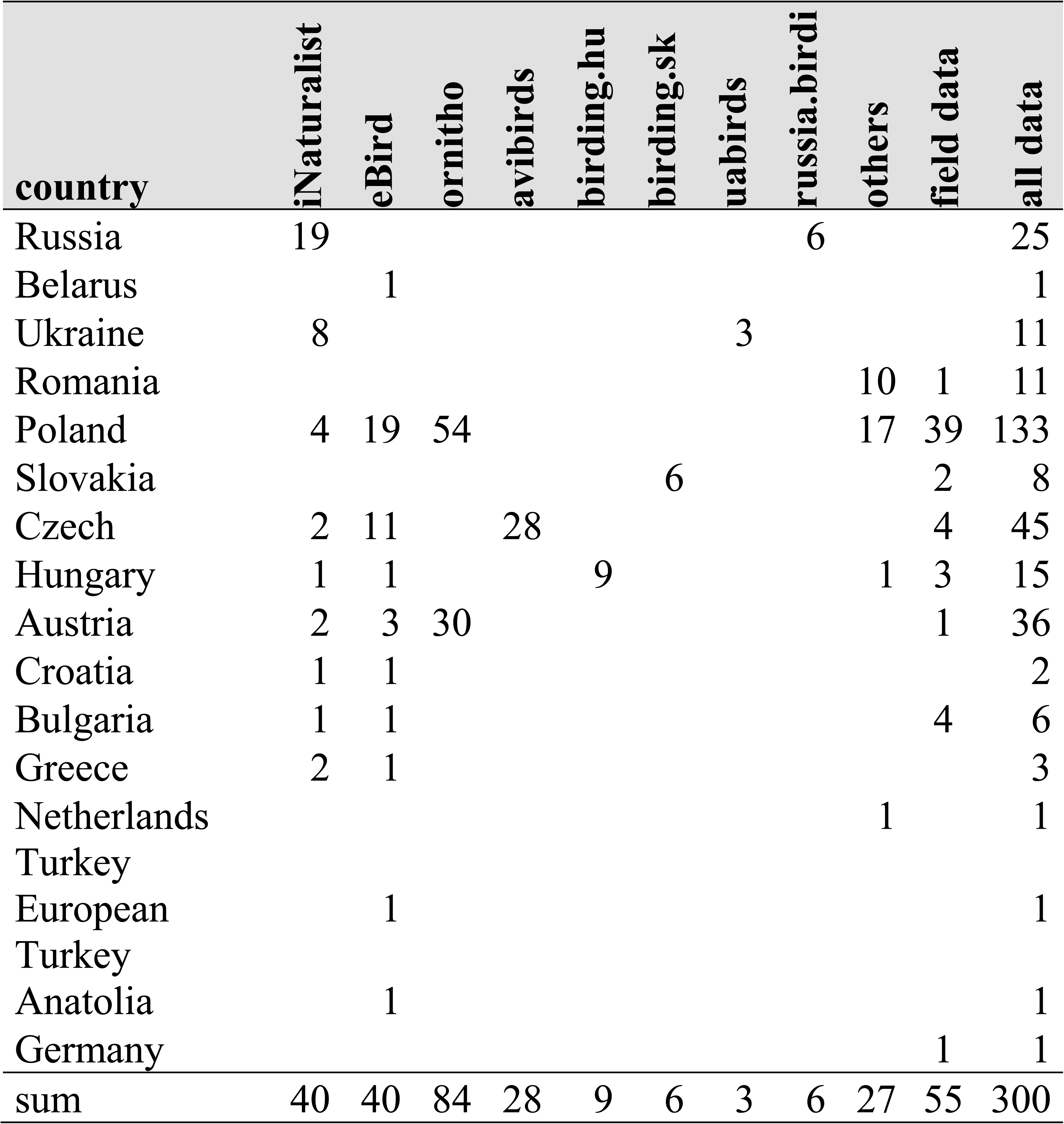
Number of Syrian x Great spotted woodpeckers hybrids in countries based on citizen science and other source of data.

| country | iNaturalist | eBird | ornitho | avibirds | birding.hu | birding.sk | uabirds | russia.birdi | others | field data | all data |
| --- | --- | --- | --- | --- | --- | --- | --- | --- | --- | --- | --- |
| Russia | 19 |  |  |  |  |  |  | 6 |  |  | 25 |
| Belarus |  | 1 |  |  |  |  |  |  |  |  | 1 |
| Ukraine | 8 |  |  |  |  |  | 3 |  |  |  | 11 |
| Romania |  |  |  |  |  |  |  |  | 10 | 1 | 11 |
| Poland | 4 | 19 | 54 |  |  |  |  |  | 17 | 39 | 133 |
| Slovakia |  |  |  |  |  | 6 |  |  |  | 2 | 8 |
| Czech | 2 | 11 |  | 28 |  |  |  |  |  | 4 | 45 |
| Hungary | 1 | 1 |  |  | 9 |  |  |  | 1 | 3 | 15 |
| Austria | 2 | 3 | 30 |  |  |  |  |  |  | 1 | 36 |
| Croatia | 1 | 1 |  |  |  |  |  |  |  |  | 2 |
| Bulgaria | 1 | 1 |  |  |  |  |  |  |  | 4 | 6 |
| Greece | 2 | 1 |  |  |  |  |  |  |  |  | 3 |
| Netherlands |  |  |  |  |  |  |  |  | 1 |  | 1 |
| Turkey |  |  |  |  |  |  |  |  |  |  |  |
| European |  | 1 |  |  |  |  |  |  |  |  | 1 |
| Turkey |  |  |  |  |  |  |  |  |  |  |  |
| Anatolia |  | 1 |  |  |  |  |  |  |  |  | 1 |
| Germany |  |  |  |  |  |  |  |  |  | 1 | 1 |
| sum | 40 | 40 | 84 | 28 | 9 | 6 | 3 | 6 | 27 | 55 | 300 |

**Table S4.**
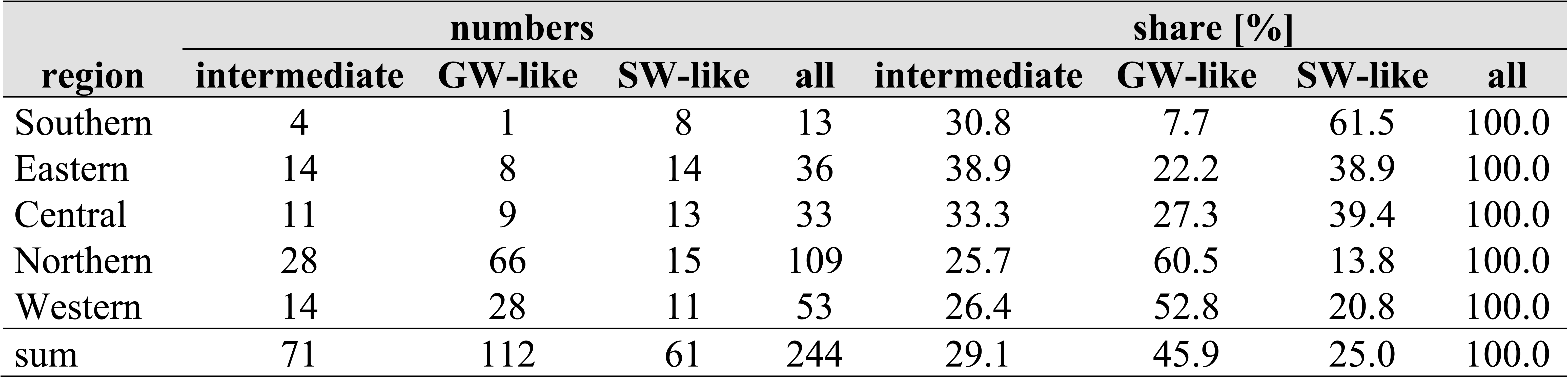
Numbers and shares of Syrian x Great spotted Woodpeckers hybrid forms in geographic regions. Great spotted Woodpecker-like (GW-like), Syrian Woodpecker-like (SW-like), intermediate.

| region | numbers |  |  |  | share [%] |  |  |  |
| --- | --- | --- | --- | --- | --- | --- | --- | --- |
|  | intermediate | GW-like | SW-like | all | intermediate | GW-like | SW-like | all |
| Southern | 4 | 1 | 8 | 13 | 30.8 | 7.7 | 61.5 | 100.0 |
| Eastern | 14 | 8 | 14 | 36 | 38.9 | 22.2 | 38.9 | 100.0 |
| Central | 11 | 9 | 13 | 33 | 33.3 | 27.3 | 39.4 | 100.0 |
| Northern | 28 | 66 | 15 | 109 | 25.7 | 60.5 | 13.8 | 100.0 |
| Western | 14 | 28 | 11 | 53 | 26.4 | 52.8 | 20.8 | 100.0 |
| sum | 71 | 112 | 61 | 244 | 29.1 | 45.9 | 25.0 | 100.0 |

**Table S5.**
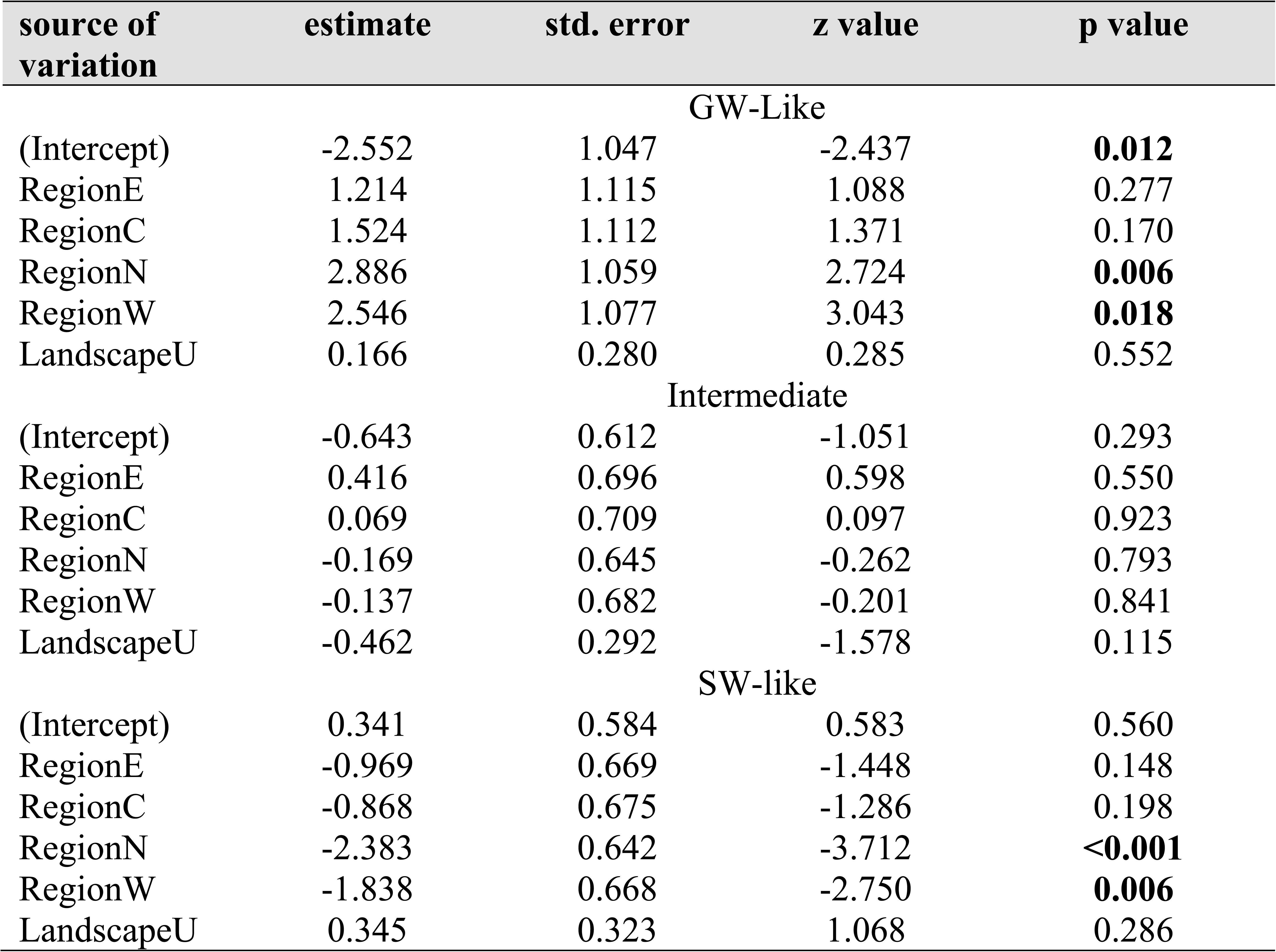
Summary of generalized of linear model (GLM) assessing the influence of region (S – shouthern, E – eastern, C – central, N – northern, W – western) and landscape type (U – urban, R – rural) on probability of Great spotted (GW)-like, intermediate and Syrian Woodpecker (SW)-like phenotypes occurrence. Probability values of significant factors (p≤0.05) are in bold.

| source of variation | estimate | std. error | z value | p value |
| --- | --- | --- | --- | --- |
| GW-Like |  |  |  |  |
| (Intercept) | -2.552 | 1.047 | -2.437 | **0.012** |
| RegionE | 1.214 | 1.115 | 1.088 | 0.277 |
| RegionC | 1.524 | 1.112 | 1.371 | 0.170 |
| RegionN | 2.886 | 1.059 | 2.724 | **0.006** |
| RegionW | 2.546 | 1.077 | 3.043 | **0.018** |
| LandscapeU | 0.166 | 0.280 | 0.285 | 0.552 |
| Intermediate |  |  |  |  |
| (Intercept) | -0.643 | 0.612 | -1.051 | 0.293 |
| RegionE | 0.416 | 0.696 | 0.598 | 0.550 |
| RegionC | 0.069 | 0.709 | 0.097 | 0.923 |
| RegionN | -0.169 | 0.645 | -0.262 | 0.793 |
| RegionW | -0.137 | 0.682 | -0.201 | 0.841 |
| LandscapeU | -0.462 | 0.292 | -1.578 | 0.115 |
| SW-like |  |  |  |  |
| (Intercept) | 0.341 | 0.584 | 0.583 | 0.560 |
| RegionE | -0.969 | 0.669 | -1.448 | 0.148 |
| RegionC | -0.868 | 0.675 | -1.286 | 0.198 |
| RegionN | -2.383 | 0.642 | -3.712 | **<0.001** |
| RegionW | -1.838 | 0.668 | -2.750 | **0.006** |
| LandscapeU | 0.345 | 0.323 | 1.068 | 0.286 |

**Table S6.**
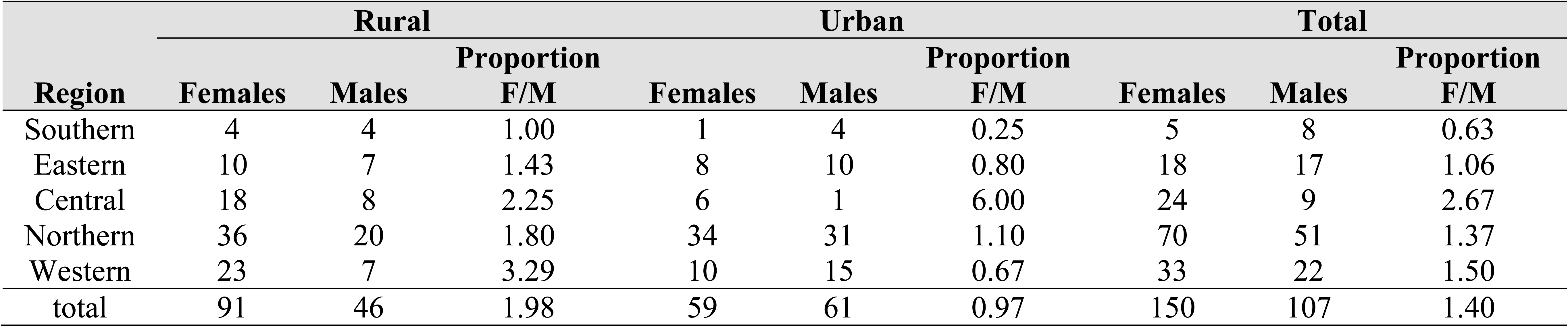
Numbers and shares of Syrian x Great spotted woodpeckers hybrid sexes in geographic regions.

**Table S7.** Summary of generalized linear model (GLM) assessing the influence of region (S – southern, E – eastern, C – central, N – northern, W – western) and landscape type (U – urban, R – rural) on Great spotted Woodpecker × Syrian Woodpecker hybrids’ sex proportion (female/male). Probability values of significant factors (p≤0.05) are in bold.

| source of variation | estimate | std. error | z value | p value |
| --- | --- | --- | --- | --- |
| (Intercept) | -2.552 | 1.047 | -2.437 | **0.004** |
| RegionE | 1.214 | 1.115 | 1.088 | 0.165 |
| RegionN | 1.524 | 1.112 | 1.371 | 0.311 |
| RegionS | 2.886 | 1.059 | 2.724 | 0.051 |
| RegionW | 2.546 | 1.077 | 3.043 | 0.393 |
| LandscapeU | 0.166 | 0.280 | 0.285 | **0.010** |

## Literature

AVIF. 2025. Galerie. https://avif.birds.cz/search/gallery?filter=%7B%22minRarityLevel%22%3A30%2C%22dateNDays%22%3A365%7D (accessed 01 August 2025)

Bakai, A., Fuchs, J., Gorman, G., Sajdak, D., Kajtoch, Ł., 2025a. A systematic review of interspecific breeding in woodpeckers. Ibis. 10.1111/ibi.70014

Bakai, A., Gorman, G., Kajtoch, Ł., 2025b. A review of co-occurrence and hybridization as neglected factors in studies of Syrian Woodpecker (*Dendrocopos syriacus*) and Great Spotted Woodpecker (*Dendrocopos major*). J. Ornithol. 10.1007/s10336-025-02298-w

Barrowclough, G.F., Groth, J.G., Bramlett, E.K., Lai, J.E., Mauck, W.M., 2017. Phylogeography and geographic variation in the Red-bellied Woodpecker (Melanerpes carolinus): characterization of mtDNA and plumage hybrid zones. Wilson J. Ornithol. 130, 671–683. 10.1676/17-070.1

Bates, D., Maechler, M., Bolker, B., Walker, S., 2015. Fitting Linear Mixed-Effects Models Using lme4. J. Stat. Softw. 67, 1–48. doi:10.18637/jss.v067.i01

Birding.hu Hungarian birdwatchers’ site. 2025. Galleries, Common birds. http://birding.hu/galeriak/gyakori_madarfajok.html (accessed 01 August 2025)

Birding Romania. 2025. Photo Galleries. https://rombird.ro/en/h/7/photo-galleries (accsessed 01 August 2025)

Birding Slovakia. 2025. Observations, Galleries. https://birding.sk/index.php/en/observations/galleries (accessed 01 August 2025)

Brooks, M.E., Kristensen, K., van Benthem, K.J., Magnusson, A., Berg, C.W., Nielsen, A., Skaug, H.J., Maechler, M., Bolker, B.M., 2017. glmmTMB Balances Speed and Flexibility Among Packages for Zero-inflated Generalised Linear Mixed Modeling. R J. 9, 378–400. 10.32614/RJ-2017-066

Dudzik, K., Polakowski, M., 2011. The cases of mixed broods and identification of Syrian Woodpecker *Dendrocopos syriacus* and Great Spotted Woodpecker *Dendrocopos major* hybrids in Poland. Chrońmy Przyr. Ojcz. 67, 254–260.

eBird, 2025. eBird: An online database of bird distribution and abundance [web application]. eBird, Cornell Lab of Ornithology, Ithaca, New York. Available: http://www.ebird.org. (Accessed: December 16, 2025).

Figarski, T., 2017. Contrasting seasonal reactions of two sibling woodpeckers to playback stimulation in urban areas—implications for inventory and monitoring of the Syrian Woodpecker. Behaviour. 154, 981–996. 10.1163/1568539X-00003452

Figarski, T., Kajtoch, Ł., 2018. Hybrids and mixed pairs of Syrian and Great Spotted Woodpeckers in urban populations. J. Ornithol. 159, 311–314. 10.1007/s10336-017-1513-0

Fitzpatrik, B.M., Shaffer, H.B., 2007. Introduction history and habitat variation explain the landscape genetics of hybrid tiger salamanders. Ecol. Appl. 17, 598–608. 10.1890/06-0369

Gorman, G., 1997. Hybridisation by Syrian Woodpeckers. Brit. Birds. 90, 578.

Gorman, G., 1999. The identification of Syrian Woodpecker. Alula. 3, 82–88.

Gorman, G., 2014. Woodpeckers of the World: The Complete Guide. A&C Black Publishers, Bloomsbury Publishing, London.

Grouw, H., Tolkmit, D., Becker, D., Weihe, F., 2021. Unusual phaeomelanised plumage: a review of similar cases in different species. Brit. Birds. 114, 629–637.

Gurgul, A., Miksza-Cybulska, A., Szmatoa, T., Semik-Gurgul, E., Jasielczuk, I., Bugno-Poniewierska, M., Figarski, T., Kajtoch L., 2019. Evaluation of genotyping by sequencing for population genetics of sibling and hybridizing birds: an example using Syrian and Great Spotted Woodpeckers. J. Ornithol. 160, 287–294. 10.1007/s10336-018-1601-9

iNaturalist contributors, iNaturalist (2025). iNaturalist Research-grade Observations. iNaturalist.org. (accessed 01 August 2025)

Kajtoch, Ł., Figarski, T., 2017. Comparative distribution of Syrian and Great Spotted Woodpeckers in different landscapes of Poland. Folia Zool. 66, 29–36. 10.25225/fozo.v66.i1.a5.2017

Kajtoch, Ł., Kusal, B., 2022. The first case of a successful brood from a double hybrid mixed pair (*Dendrocopos syriacus x Dendrocopos major* (Picidae)). Ibis. 164, 1273–1277. 10.1111/ibi.13096

Kajtoch, Ł., Kusal, B., 2023. Decline in the population of the Syrian Woodpecker *Dendrocopos syriacus* in the Krakow agglomeration. Ornis Pol. 64, 119–128. 10.12657/ornis.2023.2.3

Kajtoch, Ł., Kusal, B., Bakai, A., Pochłopień, Z., 2025. Wiślańsko-szreniawska ostoja dzięcioła białoszyjego *Dendrocopos syriacus* w województwie małopolskim. Ornis Pol. 66, 186–195.

Karlík, E., Veľký, M., 2007. Albinism in a woodpecker *Dendrocopos major/syriacus*. Trichodroma. 19, 27–48.

Keller, V., Herrando, S., Voříšek, P., Franch, M., Kipson, M., Milanesi, P., Martí, D., Anton, M., Klvanova, A., Kalyakin, M., Bauer, H., Foppen, R., 2020. European Breeding Bird Atlas 2: Distribution, Abundance and Change. European Bird Census Council & Lynx Edicions, Barcelona.

Lenth, R., 2025. _emmeans: Estimated Marginal Means, aka Least-Squares Means. doi:10.32614/CRAN.package.emmeans <10.32614/CRAN.package.emmeans>, R package version 1.11.2-8, <https://CRAN.R-project.org/package=emmeans>.

Lešo, P., Kropil, R., Kajtoch, Ł., 2019. Effects of forest management on bird assemblages in oak-dominated stands of the Western Carpathians – Refuges for rare species. For. Ecol. Manag. 453, 117620. 10.1016/j.foreco.2019.117620

Maier, M.J., 2014. DirichletReg: Dirichlet Regression for Compositional Data in R. Research Report Series / Department of Statistics and Mathematics, 125. WU Vienna University of Economics and Business, Vienna. 10.57938/ad3142d3-2fcd-4c37-aec6-8e0bd7d077e1

Melnikov, E., 2015. A female Great Spotted Woodpecker *Dendrocopos major* helps nesting pair of Syrian Woodpeckers *D. syriacus* incubate clutch and feed the nestlings. Russkii Ornitologicheskii Zhurnal Ekspress Vypusk. 1217, 4211–4214.

Michalczuk, J., 2014. Expansion of the Syrian Woodpecker *Dendrocopos syriacus* in Europe and Western Asia. Ornis Pol. 55, 149–161. 10.12657/ornis.2014.3.1

Michalczuk, J., Michalczuk, M., 2006a. Reaction on playback and density estimations of Syrian Woodpecker *Dendrocopos syriacus* in agricultural areas of SE Poland. Acta Ornithol. 41, 33–39.

Michalczuk, J., Michalczuk, M., 2006b. The usefulness of the mapping method with playback in estimation of the numbers of the Syrian Woodpecker *Dendrocopos syriacus*. Not. Ornitol. 47, 175–184.

Michalczuk, J., Michalczuk, M., 2015. Decline of the Syrian Woodpecker *Dendrocopos syriacus* population in rural landscape in SE Poland in 2004-2012. Ornis Polonica. 56, 67–75. 10.12657/ornis.2015.2.1

Michalczuk, J., Michalczuk, M., 2016. Habitat preferences of Picidae woodpeckers in the agricultural landscape of SE Poland: Is the Syrian Woodpecker *Dendrocopos syriacus* colonizing a vacant ecological niche? North-West. J. Zool. 12, 14–21.

Michalczuk, J., Michalczuk, M., 2023. The requirements of three co-existing woodpecker species (Picidae) in relation to forest features in the agricultural landscape of SE Poland. Balt. For. 29, id 561. 10.46490/BF561

Michalczuk, J., Michalczuk, M., Cymbała, R., 2011. The usefulness of various methods of monitoring the population size of the Syrian Woodpecker *Dendrocopos syriacus*. Ornis Pol. 52, 280–287.

Michalczuk, J., McDevitt, A.D., Mazgajski, T.D., Figarski, T., Ilieva, M., Bujoczek, M., Malczyk, P., Kajtoch, Ł., 2014. Tests of multiple molecular markers for the identification of Great Spotted and Syrian Woodpeckers and their hybrids. J. Ornithol. 155, 591–600. 10.1007/s10336-014-1040-1

Olszewski, A., 2007. Great Spotted Woodpeckers *Dendrocopos major* of atypical coloration in the Kampinos Forest. Not. Ornitol. 48, 210–213.

Ornitho.at, 2025. Foto and sound gallery. https://www.ornitho.at/index.php?m_id=7 (accessed 01 August 2025)

Ornitho.de, 2025. Photo and sound gallery. https://www.ornitho.de/index.php?m_id=7 (accessed 01 August 2025)

Ornitho.pl, 2025. Galleries. https://www.ornitho.pl/index.php?m_id=7 (accessed 01 August 2025)

Ottenburghs, J., Nicolaï, M.P., 2024. Hybridization constrains the evolution of mimicry complexes in woodpeckers. J. Avian Biol. 2024, e03228. 10.1111/jav.03228

Posit team, 2024. RStudio: Integrated development environment for R. Posit Software, PBC, Boston, MA. URL http://www.posit.co/.

Prum, R.O., Brev, J.S., Dornburg, A., Field, D.J., Townsend, J.P., Lemmon, E.M., Lemmon, A.R., 2015. A comprehensive phylogeny of birds (Aves) using targeted next-generation DNA sequencing. Nature. 526, 569–573. 10.1038/nature15697

R Core Team, 2025. R: A language and environment for statistical computing [software]. R Foundation for Statistical Computing, Vienna, Austria. URL https://www.R-project.org/.

QGIS.org, 2025. QGIS Geographic Information System [software]. QGIS Association. http://www.qgis.org

Seneviratne, S.S., Davidson, P., Martin, K., Irwin, D.E., 2016. Low levels of hybridization across two contact zones among three species of woodpeckers (Sphyrapicus sapsuckers). J. Avian Biol. 47, 887–898. 10.1111/jav.00946

Shakya, S.B., Fuchs, J., Pons, J.M., Sheldon, F.H., 2017. Tapping the woodpecker tree for evolutionary insight. Mol. Phyl. Evol. 116, 182–191. 10.1016/j.ympev.2017.09.005

Short, L.L., 1982. Woodpeckers of the World. Monograph Series 4. Delaware Museum of Natural History, Greenville, DE, USA.

Sullivan, B.L., Wood, C.L., Iliff, M.J., Bonney, R.E., Fink, D., Kelling, S., 2009. eBird: a citizen-based bird observation network in the biological sciences. Biol. Conserv. 142, 2282–2292.

Uabirds org Ukrainian Birdwatching Community. 2025. Syrian woodpecker, Gallery. https://uabirds.org/v2taxgal.php?s=1451&l=en&p=0 (accessed 01 August 2025)

Ukrainian Biodiversity Information Network. 2025. Gallery. https://ukrbin.com/index.php?action=taxa (accessed 01 August 2025)

van Grouw, H., Tolkmit, D., Becker, D., Weihe, F., 2021. Unusual phaeomelanised plumage: a review of similar cases in different species. British Birds. 114: 629–637.

Venables, W.N., Ripley, B.D., 2002. Modern Applied Statistics with S. Fourth Edition. Springer, New York. ISBN 0-387-95457-0

Wickham, H., 2016. ggplot2: Elegant Graphics for Data Analysis. Springer-Verlag New York.

Winkler, H., 1971. Die artliche Isolation des Blutspechts *Picoides (Dendrocopos) syriacus*. Egretta. 14: 1–20.

Winkler, H., Christie, D.A., Nurney, D., 1995. Woodpeckers, a Guide to the Woodpeckers of the World. Houghton Mifflin Company, Ne York.

Venables, W.N., Ripley, B.D., 2002. Modern Applied Statistics with S, fourth ed. Springer, New York.

